# DNMT/G9a Complex Inhibition Uncovers Epigenetic Vulnerabilities and Induces IFN-Response in Acute Myeloid Leukemia

**DOI:** 10.1101/2024.12.11.627891

**Authors:** Angelique Schönefeld, Costanza Zanetti, Eric C. Schmitt, Kevin Woods, Ashley Westerback, Sidney M. Mitsch, Frank Rühle, Simge Kelekci, Lisa Fol, Viral Shah, Miriam Sanchez-Saez, Sophia Thevissen, Alexander Schäffer, Ulrike Regina Schmits, Daniel Sasca, Thomas Kindler, Catherine Wölfel, Clarissa Feuerstein-Akgoz, Clarissa Gerhäuser, Pavlo Lutsik, Jennifer Rivière, Judith S Hecker, Katharina S Götze, Uwe Platzbecker, Hakim Echchannaoui, Matthias Theobald, Christoph Plass, Jan Padeken, Hind Medyouf, Borhane Guezguez

## Abstract

Epigenetic dysregulation is a hallmark of Acute Myeloid Leukemia (AML), with mutations in DNA Methyltransferases (e.g., DNMT3A) being frequent and promising therapeutic targets. DNMTs form complexes with Histone Methyltransferases (HMTs), driving gene silencing loop via chromatin methylation crosstalk. However, potential connections between this DNMTs/HMTs cooperative activity and oncogenic requirements across the AML mutational spectrum remain poorly understood. Here, we demonstrate that AMLs carrying DNMT3A and Nucleophosmin (NPM1) mutations exhibit a specific epigenetic vulnerability toward a complex formed by DNMTs and G9a, a specific histone H3 Lysine 9 Methyltransferase (H3K9-HMT). Dual inhibition of DNMT/G9a restores differentiation, reduces tumor growth, and spares healthy progenitors compared to standard hypomethylating agents. Mechanistically, DNMT/G9a regulates NPM1 stability, inhibits HOXA9/MEIS1 activity, and triggers interferons (IFN) response via viral mimicry pathways by modulating hypermethylated retrotransposons. Collectively, our data unravel specific epigenetic vulnerabilities within the complex AML mutational landscape and provide a compelling rationale for the design of personalized epigenetic therapies with enhanced efficacy and safer clinical outcomes.

## Introduction

Acute myeloid leukemia (AML) is a genetically diverse group of highly aggressive hematopoietic malignancies characterized by myeloid maturation blockade, rapid blast accumulation, and poor prognosis ^1,2^. Despite mutational heterogeneity, epigenetic dysregulation represents a common hallmark in AML, providing potential pharmacological targets for therapeutic intervention ^3^. Among these, mutations of the DNA methyltransferase 3A (DNMT3A), a de novo DNA methyltransferase in the DNA methylation pathway, are frequently detected in AML, resulting in epigenetic deregulation of key biological pathways ^4,5^. Therefore, the reversal of aberrant DNA methylation patterns with hypomethylating agents (HMA, cytidine analogs such as Decitabine or Azacytidine) has shown clinical benefit in AML patients and has recently been approved for combined chemotherapy with the BCL2 inhibitor Venetoclax, but their long-term efficacy to improve overall survival remains limited ^6^. More importantly, primary drug resistance and disease relapse remain persistent problems for treatment with HMA and other DNMT inhibitors, prompting a more comprehensive understanding of the oncogenic transcriptional programs controlled by aberrant DNA methylation in AML.

Recent evidence from large meta-analyses of genome-wide profiling studies found a strong correlation between DNA methylation and histone methylation, specifically histone H3 Lysine 9 (H3K9) methylation, a repressive chromatin mark associated with gene silencing and constitutive heterochromatin formation ^7–9^. Moreover, aberrant H3K9 methylation levels have also been linked to AML etiology and patient survival outcome ^10,11^. H3K9 can be mono-(H3K9me1), di-(H3K9me2), or tri-(H3K9me3) methylated, which is catalyzed by specific histone methyltransferases (H3K9-HMT) of the SET-containing SUV39 protein family: SUV39H1, SUV39H2, SETDB1 for H3K9me3 and G9a/GLP (EHMT2/EHMT1) for H3K9me2 ^12^. Even though H3K9me3 was found to be the most prominent epigenetic dysregulation of heterochromatin in AML ^10,13^, H3K9me2 levels and G9a/GLP activity were specifically found to be involved in repression of multiple genes associated with myeloid leukemogenesis and are hypothesized to promote chromosome instability and silencing of tumor suppressors and retrotransposons ^11,14–16^. Furthermore, recent molecular studies have uncovered that H3K9me2 methylation is physically dependent on DNA methylation via direct interaction of DNMT with G9a/GLP and their associated factors into a molecular complex to ensure a constant gene silencing loop at heterochromatin regions ^17–19^. Finally, previous pharmacological and genetic targeting of G9a/GLP was shown to be therapeutically relevant to inhibit tumor growth in a mouse model and human AML cell lines but did not impact overall DNA methylation and gene silencing ^20–22^. Therefore, further understanding of the mechanisms underlying the oncogenic dependencies of the aberrant activity of DNMT/H3K9-HMT complex in AML would be potentially beneficial for more efficient epigenetic treatments in the future.

Here we demonstrate a specific DNMT/G9a epigenetic vulnerability of an AML subgroup carrying mutations for Nucleophosmin (NPM1) and DNMT3A, which are known to account for approximately 30% of all AML cases^23^. At the molecular level, we found that aberrant expression of DNMT/G9a complex in AML promotes the neoplastic activity of the transcriptional complex HOXA9/MEIS1 and maintains NPM1 instability. Pharmacological targeting of DNMT/G9a in vitro and in vivo can reverse these aberrant genetic programs by restoring terminal myelo-monocytic differentiation and inducing DNA demethylation at specific enhancer regions with no toxic side-effects on healthy cells. More strikingly, we determine a universal role of DNMT/G9a as safeguards of viral interferon pathway activation through silencing of specific retrotransposons. Collectively, our findings provide a rationale for targeted therapy of DNMT/G9a in NPM1/DNMT3A-mutated AMLs, and more broadly hold promises in distinguishing specific epigenetic vulnerabilities within the complex AML mutational landscape for tailored personalized medicine.

## Results

### G9a and DNMT molecular profiles are mutually dysregulated in NPM1c AML

Dysregulation of G9a and its oncogenic role have been reported in a variety of cancers including AML ^24^. However, our understanding of the molecular implications of G9a and its association with aberrant DNA methylation in driving leukemogenesis remains largely unknown. Using a molecular data mining approach, we performed an in-depth analysis of G9a gene expression and its clinical significance, potential molecular functions, and regulatory networks in three large-scale public RNA-sequencing (RNA-seq) data sets from AML patients (BeatAML = 698, TCGA-LAML = 179 and MILE = 542). We found that the G9a gene transcript (*EHMT2*) is highly up-regulated in AML compared to healthy bone marrow cells **(Figure 1A)**, including other myeloid malignancies such as MDS and CML **(supplementary Figure S1A and S1B)**. Within the AML molecular landscape, elevated levels of *EHMT2/G9a* expression are evident in subtypes with normal karyotype (NK) and those harboring PML/RARα fusion gene (APL) and t(8;21) translocation **(Figure 1B)**, suggesting a possible association with specific AML mutation profiles. Next, we investigated *EHMT2/G9a* expression in AML for its biological and prognostic significance according to the European LeukemiaNet (ELN) 2017 risk classification ^2^. We found higher expression of *EHMT2/G9a* with favorable and intermediate AML subgroups **(Figure 1C)**, which is in accordance with reported prognosis for AML-NK and APL subtypes ^2^.

**Figure 1.**
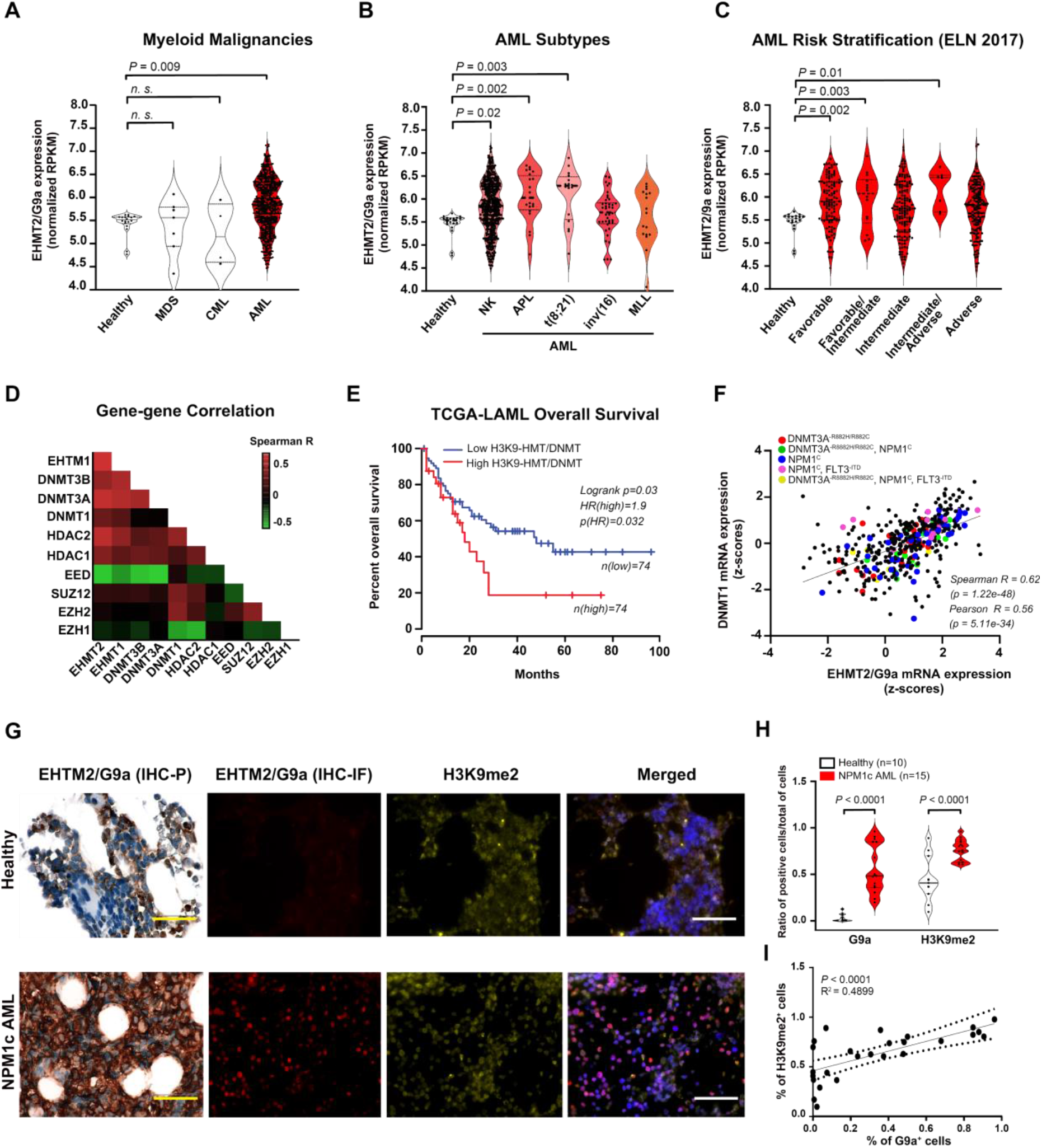
*G9*a and DNMT co-expression in AML and their association with patient survival and genetic subgroups. **(A-C)** *EHMT2 (G9a)* mRNA expression (normalized RPKM) according to **(A)** hematological disease subtypes, **(B)** cytogenetic/WHO-Fusion AML subtypes and **(C)** AML prognostic risk categories (ELN2017) from the BeatAML study (n=707 samples). Within each Violin plot, a circle represents one patient value, dotted lines delimit first and third quartiles and black bars indicate the median. *P*-value statistical significance was assessed by one-way ANOVA with Dunnett’s correction for multiple comparisons (each condition vs. healthy bone marrow). The non-indicated comparisons were not significant (*P* > 0.05). **(D)** Spearman correlation heat map between the relative mRNA levels of indicated genes in BeatAML study. The red color indicates a positive correlation, the green color indicates a negative correlation and the black color indicates no correlation. Spearman R values were determined from two-tailed Pearson’s correlation test. **(E)** Kaplan–Meier curve representing overall survival analysis of AML patients from the TCGA-LAML study (*n* = 148) with combined mRNA expression of H3K9-HMT/DNMT gene set above (high = 74) and below the median (low =74). Statistical significance for *P*-value and hazard ratio (HR) was assessed by log-rank and Cox test. **(F)** Correlation of the mRNA expression (z-scores) of EHTM2 and DNMT1 in AML patient-derived bone marrow cells (BeatAML study, n=707 samples). Each circle represents a patient value, colored circles indicate AML samples with specific mutational profiles (as indicated on the graph). Two-tailed Pearson’s correlation test was performed for statistical significance. Spearman’s and Pearson’s correlation coefficients (*R*) and corresponding *P* values are shown. **(G)** Representative bone marrow sections of healthy donors and NPM1c AML patients. EHMT2/G9a positive signal is assessed by Immunohistochemistry staining with specific antibody and revealed by chemiluminescence (HRP) and immunofluorescence for G9a (red), H3K9me2 marks (yellow) and nuclei (DAPI). Images represent x40 magnification. **(H)** Violin graphs represent ratio of positive G9a and H3K9m2 cells in healthy (n=10) and NPM1c AML (n=15) bone marrow sections. Within each Violin plot, a circle represents one patient value, dotted lines delimit first and third quartiles and black bars indicate the median. Two-way ANOVA test was performed for *p-*value statistical significance. **(I)** Correlation between the percentage of detected G9a-positive cells and H3K9me2-positive cells in AML bone marrow sections. Simple linear regression test was performed for statistical significance. Coefficient of determination (*R*^*2*^) and *P* value are shown.

Previous evidence demonstrated an interdependence between G9a and DNMTs in a common molecular complex to ensure gene silencing ^18^, which we confirmed in the available datasets by functional protein-protein network String analysis **(supplementary Figure S1C)**. Furthermore, to explore the relationship between G9a and other chromatin repressive regulators, we performed a co-expression matrix analysis from BeatAML RNA-seq data of 10 epigenetic modulators previously shown to recruit G9a in a repressive multicomplex ^25^. As expected, we demonstrated a strong correlation between *EHMT2/G9a* and its heterodimer G9-like protein (*GLP/EHMT1*), followed by DNMT family (*1/3A/3B*) and core members of the histone deacetylases family (*HDAC1* and *HDAC2*), but not those of the Polycomb repressive complex 2 (PRC2) including *EZH1, EZH2, EED*, and *SUZ12* **(Figure 1D)**. By further exploring these correlations toward prevalent gene clusters in both the BeatAML and TCGA-LAML datasets, we identified a distinct set of genes including *EHMT2, EHMT1, DNMT3A*, and *DNMT3B* **(supplementary Figure S1D)**. High EHMT/DNMT expression was strongly associated with unfavorable overall survival in the TCGA-LAML study **(Figure 1E)**. Of note, expression levels of each individual gene from the EHMT/DNMT signature did not have a prognostic impact on AML survival, except for DNMT3A **(supplementary Figure S1E)**. Next, we extended our co-expression matrix analysis to assess potential associations of the EHMT/DNMT gene set with specific AML subgroups based on recurrent mutations according to the WHO classification ^26^. Within the BeatAML cohort, we found a strong positive correlation between the expression of DNMT1/G9a and AML samples harboring mutations in the NPM1 gene that generate an aberrant cytoplasmic protein (hereafter referred to as NPM1c) (Spearman R= 0.62, p < 0.0001, **Figure 1F**). NPM1c AMLs define a well-known AML subset with singular clinical characteristics ^27,28^. In addition, most of these NPM1c AML samples correlating with DNMT1/G9a **(Figure 1F)** were found to harbor known co-occurring mutations for DNMT3A (mostly somatic R882H/R882C alterations) ^29^ and FMS-related tyrosine kinase 3-internal tandem duplication (FLT3-ITD) ^30^, which overall account for close to 35% of all AML cases ^23^. Therefore, we conclude that NPM1c AMLs likely exhibit dysregulated G9a and H3K9me2 molecular profiles.

We next validated if G9a overexpression is associated with enhanced di-methylation of H3K9 (H3K9me2) using immunohistochemical analyses of bone marrow biopsies from healthy (n=10) and NPM1c AML patients (n=15), **(Figure 1G and Supplementary Table S1)**. In comparison to healthy cases, we found that levels of G9a protein and the histone mark H3K9me2 were both upregulated in NPM1c AML **(Figure 1H)**, with a substantial overlapping in their staining patterns (R2 = 0.49, p < 0.0001, **Figure 1I**). Collectively, these results indicate an association of G9a and DNMT overexpression in NPM1c AMLs that is specific for a subgroup with DNMT3A mutations.

### Dual DNMT/G9a inhibition demonstrates potent efficacy in NPM1c AML

The molecular data analysis presented so far suggested the possibility to target the epigenetic activity of the DNMT/G9a complex in NPM1c AML harboring DNMT3A mutations (with or without FLT3-ITD). Consistently, the functional genomic data of the BeatAML study demonstrated that NPM1c AMLs were most sensitive to DNMT inhibition with (Azacytidine, **supplementary Figure S2A**), in comparison to treatment with other chemotherapeutic agents such as the DNA Polymerase inhibitor (Cytarabine, **supplementary Figure S2B**), the BCL2 inhibitor (Venetoclax, **supplementary Figure S2C**), or other epigenetic regulators such as bromodomain (OTX-015, **supplementary Figure S2D**) or histone deacetylase (HDAC) inhibitors (Panobinostat, **supplementary Figure S2E**).

We experimentally evaluated the synergistic pharmacological inhibition by measuring the cytotoxic dose response (IC_50_) of the DNMT/G9a complex dual inhibitor (CM-272: IC_50_ = 241 nM) in DNMT3A^R882C^ NPM1c AML cells (OCI-AML3 cell line) compared to treatment with a standard HMA (Decitabine or DAC: IC_50_ = 746 nM), a stand-alone G9a inhibitor (UNC-0638: IC_50_ = 2.5 μM) and a specific DNMT inhibitor (SGI-1027: IC_50_ = 628 nM) **(Figure 2A)**. To determine if the DNMT1/G9a complex is a valid therapeutic target in other AML subtypes, we assessed the cytotoxicity of the dual inhibitor in a representative panel of AML cell lines with different mutational backgrounds. We observed that CM-272 was most potent, at concentrations ≤ 200 nM, in AML cell lines harboring primarily DNMT3A or NPM1 mutations (OCI-AML3: IC_50_ = 255 nM, OCI-AML2: IC_50_ = 114 nM) or FLT3-ITD with MLL rearrangements (MOLM-13: IC_50_ = 273 nM). In contrast, the potency was reduced (IC_50_ ≥ 500 nM) in cell lines of a myelodysplasia/erythroleukemia origin (F-36P: IC_50_ = 544 nM, MDS-L: IC_50_ = 482 nM) or harboring other recurrent mutations (Kasumi-1: IC_50_ = 469 nM, KG1a: IC_50_ = 7.15 μM). Therefore, these in vitro results are consistent with the molecular finding presented in **Figure 1**.

**Figure 2.**
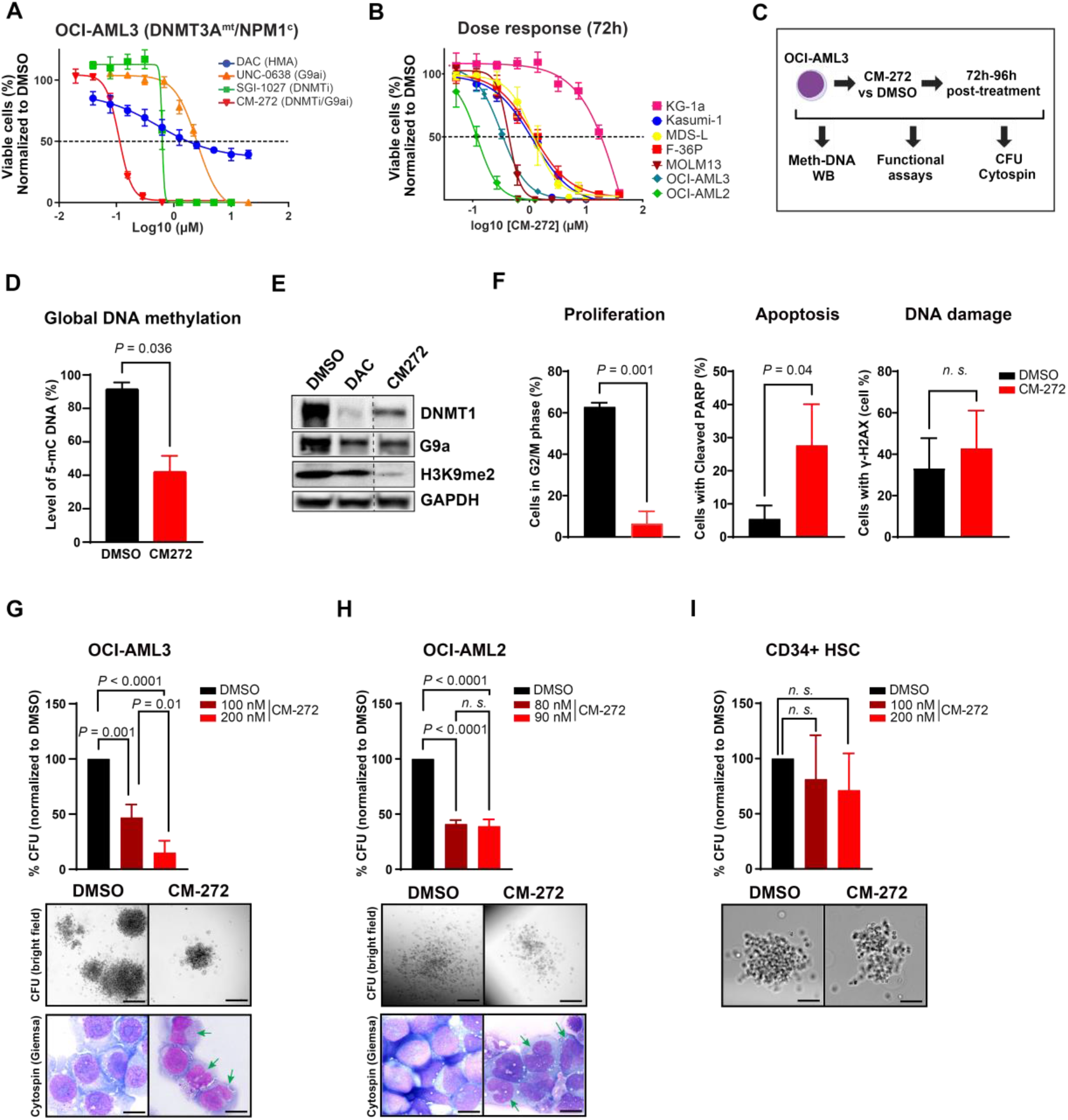
Impact of dual targeting DNMT/G9a on patient-derived AML cell lines growth. **(A)** Dose-response curves from cell-viability assays of OCI-AML3 cells after 72h treatment with Decitabine (DAC), CM-272 and SGI-1027. Drug treatments were normalized to DMSO values and IC-50s were determined from three independent experiments performed in technical triplicate. **(B)** Dose-response curves of different MDS/AML cell lines after 72h treatment with nine different concentrations of CM-272 in a single dose. Drug treatments were normalized to DMSO values and IC-50s were determined from three independent experiments performed in technical triplicate. **(C)** Experimental design of CM-272 treatment and functional assays performed at 72-96h post-treatment. **(D)** Level of DNA Methylation (% of 5-mC) in OCI-AML3 cells after 200nM CM-272 (single dose, 96h) compared with the DMSO control. Bar graph represents fold change expression [mean ± standard deviation (s.d)] of 3 independent experiments performed in technical triplicate. Welch’s *t*-test was performed for *p-*value statistical significance. **(E)** Immunoblot analysis of indicated protein levels in OCI-AML3 upon treatment with 500nM DAC (daily, 72h) and 200nM CM-272 (single dose, 72h). One representative blot of 2 independent experiments is shown. **(F)** Percentage of Cell cycle (% of G2/M-phase), Apoptosis (% of Cleaved PARP) and DNA damage (% of *γ*-H2AX) of OCI-AML3 cells after 200nM CM-272 (single dose, 72h) compared with the DMSO control. Bar graph represents mean ± s.d of 3 independent experiments performed in technical triplicate. Welch’s *t*-test was performed for *p-*value statistical significance. **(G)** Colonies outgrowth of OCI-AML3, **(H)** OCI-AML2 and **(I)** CD34+ HSC cells after CM-272 treatment at different indicated concentrations. **(G-I)** CFU assays were carried out in technical duplicates and scored after 10-12 days incubation in methylcellulose. Data are presented as colonies percentage normalized to DMSO control ± s.d. One-way ANOVA was performed for *p-*value statistical. Micrographs of representative CFUs and Giemsa-stained cytospins were taken at x60 amplification.

To further support the finding that AML harboring DNMT3A and/or NPM1 mutations are among the most sensitive to therapeutic targeting of DNMT/G9a complex by CM-272, we accessed data of a CRISPR/Cas9 dropout screen in AML cell lines (n = 24) (https://depmap.org/portal/) ^31^. Based on gene effect scores (Chronos), both inactivation of DNMT1 and G9a were the most deleterious to AML cell lines compared to DNMT3A and GLP/EHMT1 **(supplementary Figure S3A)**. More specifically, OCI-AML3 and OCI-AML2 showed the greatest susceptibility to gene inactivation of both DNMT1 and G9a **(supplementary Figure S3B)**. This is consistent with previous chemical findings that CM-272 is more selective toward both inhibition of DNMT1 and G9a ^32^.

Accordingly, we focused our downstream functional experiments by treating OCI-AML3 cells (DNMT3A^R882C^ NPM1c) with CM-272 for around 72 to 96 h (2.5 doubling times) **(Figure 2C)**. We observed a significant decrease in global DNA methylation (5-methylcytosine: 5meC, **Figure 2D**) and H3K9me2 levels **(Figure 2E)**. CM-272 treatment substantially decreases the protein levels of both DNMT1 and G9a, possibly due to proteosomal activation as reported previously ^33^. Inhibition of DNMT/G9a activity by CM-272 induced rapid cell cycle arrest and apoptosis without DNA damage, thereby exerting a less deleterious genotoxic effect **(Figure 2F)** than observed with decitabine or other HMA formulations ^34^. Sensitivity to CM-272 was also apparent in clonogenic progenitor assays, where colony growth gradually decreased with increasing drug concentrations in OCI-AML3 and OC-AML2 (between 80 to 200 nM, **Figure 2G and H**). Importantly, at these doses (100 and 200 nM), CM-272 did not display any effects on healthy hematopoietic progenitor cells (CD34^+^ UCB, **Figure 2I**). To a lesser extent, erythroleukemia/myelodysplastic cell lines (MDS-L, F-36P) also showed sensitivity to CM272 requiring higher drug concentrations (700 and 1000 nM, **supplementary Figure S3C-D**).

Overall, these data demonstrate a strong DNMT/G9a-dependent epigenetic vulnerability that preferentially sensitizes DNMT3Amut NPM1c AMLs toward CM-272, while preserving healthy cells.

### DNMT/G9a targeting reveals dependencies on transcriptional and molecular processes in NPM1c AML

To assess global transcriptional changes after DNMT/G9a complex inhibition in DNMT3Amut NPM1c AML cells, we performed RNA-seq analysis after treating OCI-AML3 cells with 200 nM CM-272 for 72 h. We found more genes to be upregulated (947) than downregulated (398) (by at least two-fold and adjusted P ≤ 0.01). Many of the highest upregulated genes were associated with myelo-monocytic differentiation **(Figure 3A, supplementary Figure S4A)**, including lineage-specific cell surface markers (*CD14, CD163, FCGR3A/CD16, CSF1R/CD115, MARCO*) and specific transcription factors (*MAFB, EGR2*) ^35,36^. As anticipated, known target genes of the oncogenic HOXA9/MEIS1 complex (*MEIS1, FLT3, BCL2*) ^37^ as well as markers of immature myeloid cell lineage (*ELANE, CTSG, PRTN3*) were found to be highly suppressed ^38,39^. Importantly, cell cycle/apoptosis regulators (*CDKN2A, CDKN2B*), which are suppressed by HOXA9/MEIS1 were strongly re-expressed in response to CM-272 **(Figure 3A, supplementary Figure S4A)**, which is consistent with the observed apoptotic response **(Figure 2F)**. Extended gene set enrichment analysis (GSEA) revealed specific dysregulated pathways **(Figure 3B)** with up-regulation of gene signatures associated with monocytic lineage commitment, repression of HOXA9/MEIS1 targets and HMA responsive targets **(supplementary Figure S4B)**, whereas gene signatures associated with oncogenic activity of MYC and EZH2 were depleted in response to CM-272 **(supplementary Figure S4C)**. Flow cytometry analysis further revealed an increased percentage of cells with CD14 and CD11b cell surface expression in CM272-treated OCI-AML3 cells **(Figure 3C)** along with decreased expression of FLT3 and MEIS1 as assessed by q-PCR **(Figure 3D)**.

**Figure 3.**
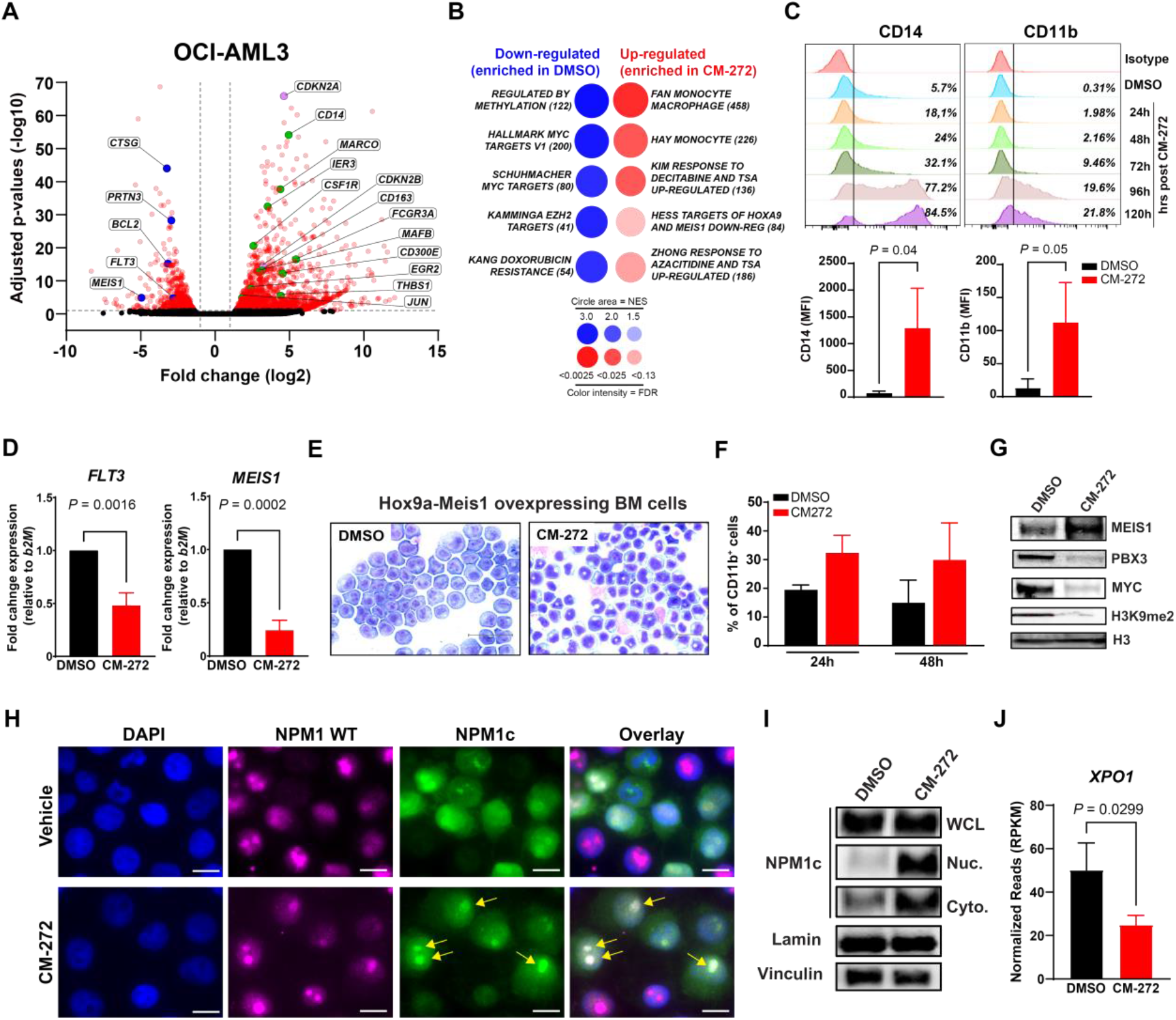
Molecular changes in in DNMT3A^mut^ NPM1^c^ AML cells upon inhibition of DNMT/G9a complex. **(A)** Volcano plots of RNA-seq data obtained from OCI-AML3 treated with CM-272 (200nM). Genes involved in Myelo-monocytic differentiation, cell cycle regulation and HOXA9 targets are labeled in green, magenta and blue respectively. **(B)** BubbleGUM gene set enrichment analysis (GSEA) map for most significant pathways of OCI-AML3 RNA-seq established from MsigDB database. For each gene set, origin of the data set and number of upregulated (red, CM272 treated) and downregulated (blue, DMSO control treated) genes are indicated. The panel summarizes the normalized enrichment score (NES) and FDR parameters. **(C)** CD14 and CD11b protein cell surface expression in OCI-AML3 cells at different time points after treatment with CM-272 (200nM, single dose), as assessed by flow cytometry. One representative histogram of 3 independent experiments is shown. The colored numbers in the flow histograms indicate the percentage of CD14 and CD11b, respectively. Bar graph represents mean Fluorescence Intensity (MFI) [mean ± standard deviation (s.d)] of 3 independent experiments performed in technical triplicate. Welch’s *t*-test was performed for *p-*value statistical significance. **(D)** Gene expression of HOXA9 gene targets (*FLT3, MEIS1*) in OCI-AML3 post-CM272 treatment as assessed by q-PCR. Bar graph represents fold change expression [mean ± standard deviation (s.d)] of 3 independent experiments performed in technical triplicate. Welch’s *t*-test was performed for *p-*value statistical significance. **(E-G)** Data related to Hoxa9-Meis1 overexpressing bone marrow cells post treatment with CM-272 (200nM, single dose). **(E)** Representative Giemsa-stained cytospins post 72h treatment and taken at x20 amplification. **(F)** CD11b expression at 24h and 48h. Bar graph represents percentage of FACS-gated positive cells [mean ± standard deviation (s.d)] of 3 independent experiments performed in technical triplicate. **(G)** Immunoblot analysis of MEIS1, PBX3, MYC and H3K9me2 protein levels upon treatment with 200nM CM-272 (single dose, 72h). One representative blot of 2 independent experiments is shown. **(H)** Immunofluorescence microscopy analysis of NPM1 localization in OCI-AML3 cells after treatment with CM-272 for 48 hours. NPM1 was stained with specific antibodies recognizing wild type form (NPM1-WT, red) and mutant form (NPM1c, green) and nuclei were stained with DAPI (blue). Images represent x100 magnification and arrows indicate nucleoli localization. **(I)** Immunoblot analysis of mutant NPM1c protein levels after sub-cellular fractionation (Whole lysate, nuclear and cytoplasmic) in OCI-AML3 upon treatment with 200nM CM-272 (single dose, 72h). One representative blot of 2 independent experiments is shown. **(J)** Gene expression of NPM1 co-factor (*XPO-1*) in OCI-AML3 post-CM272 treatment (RNA seq, three biological replicates for each condition). Data are presented as normalized reads ± s.d.

Previous studies demonstrated that maintenance of leukemia state by the Hoxa9/Meis1 complex is dependent on the recruitment of G9a for transcriptional repression ^20^. Although levels of HOX genes were unaffected after treatment **(supplementary Figure S4D)**, we hypothesize a possible direct role of CM-272 by targeting the HOXA9/G9a interaction. In order to avoid possible indirect or hidden effects of NPM1c, we tested CM-272 efficacy on Hoxa9/Meis1-transformed immature murine bone marrow cells ^40^. Following treatment, CD11b expression and cytological alterations confirmed the induction of myeloid differentiation **(Figure 3E-F)**. Additionally, reduction of protein levels of PBX3 and MYC **(Figure 3G)**, both known molecular targets of HOXA9 ^41,42^; indicated that CM-272 directly dysregulates the HOXA9/MEIS1 complex to restore myeloid differentiation.

In parallel to the perturbation effects on the chromatin complex, we sought to test if CM-272 impacts NPM1c subcellular distribution, which is also critical for leukemia maintenance. Fluorescent antigen labeling-based microscopy of OCI-AML3 treated with CM-272 demonstrated a substantial relocalization of NPM1c in the nucleus with a specific accumulation in the nucleoli **(Figure 3H)**. In parallel, Western-blot analysis from subcellular fractionation confirmed these findings **(Figure 3I)** along with a decrease in *XPO-1* gene expression **(Figure 3J)**, known to drive NPM1c-dependent aberrant *HOX* transcription.

Taken together, these data indicate that dual inhibition of DNMT/G9a in AML by CM-272 exert multifaceted effects, including re-expression of specific H3K9me2-silenced genes, a direct reversal of a HOXA9/MEIS1-orchestrated leukemogenic gene expression program and the indirect restoration of NPM1 nuclear function, ultimately resulting in myelo-monocytic differentiation and apoptotic induction.

### DNMT/G9a complex regulates a ‘core’ transcriptional program in AML

Due to the similar phenotypic consequences following CM-272 treatment in AML cell lines **(Figure 2B)**, we hypothesize that dysregulation of a common ‘core’ transcriptional program may have mediated these similar effects. To test this hypothesis, we analyzed the changes in gene expression following CM-272 treatment of sensitive (OCI-AML3) and less sensitive (MDS-L and F-36P) AML cell lines **(Figure 2 and supplementary Figure 3)**. Compared to OCI-AML3 (947 genes), only a relatively small number of genes showed a significant change in expression in MDS-L (143 genes) and F-36P (328 genes), with the majority of genes unchanged except for essential erythro-myeloid differentiation genes **(supplementary Figure S5A-B)**. This specific decrease corroborated the specific epigenetic vulnerability of NPM1c AML to DNMT/G9a inhibition. The degree of overlap of the transcriptional changes between the three cell lines was very low, resulting in only 22 commonly up-regulated genes after CM-272 treatment **(Figure 4A)**. Notably, an up-regulated set of 13-genes was identified **(Figure 4B)** that includes key cytokines and regulators primarily involved in type I interferon (IFN) signaling pathway (*IRF7, GBP1, CCL3*), innate immune response (*THBS1, CCR2, CCL2*) and defense response to virus (*OASL, IFI6, IFI27*) ^43,44^. Among the 13-genes ‘core’, *OASL* and *IFI6* were among the highest expressed of the anti-viral regulators in all three cell lines **(Figure 4C)** and are known to target different stages of the viral replication cycle by preventing the formation of virus-induced endoplasmic reticulum membrane invaginations ^45^.

**Figure 4.**
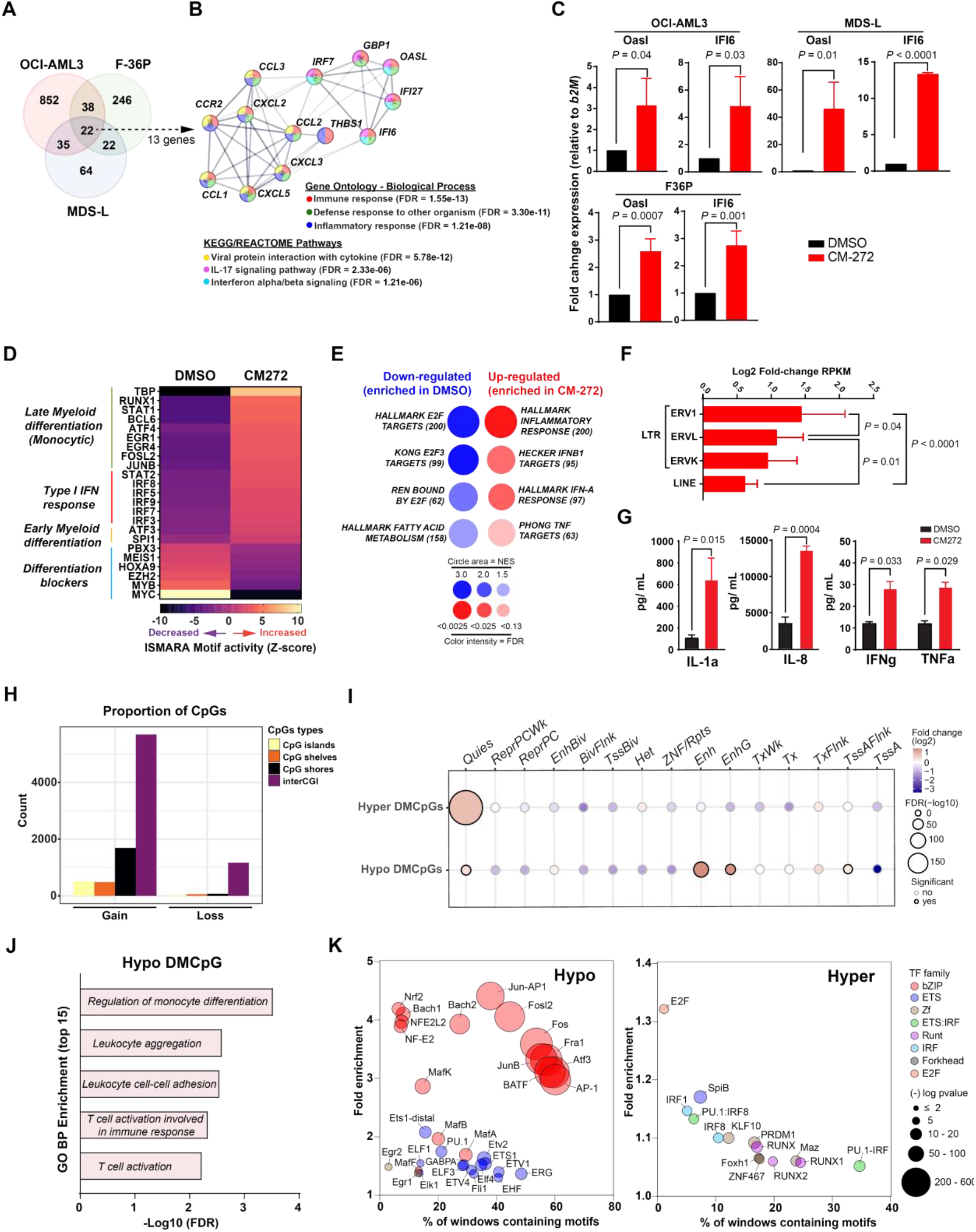
DNA methylation changes in AML cells upon inhibition of DNMT/G9a complex. **(A)** Venn diagram showing up-regulated genes identified by RNA-seq (more than two-fold increase; adjusted *P* < 0.05), in OCI-AML3, *MDS-L*, and F36P cells after CM-272 treatment compared with the DMSO control (3 biological replicates for each cell type and condition). **(B)** Mapping view of relevant functional protein association network of *1*3-genes signature generated using STRING database; stronger associations are represented by thicker lines and gene products are highlighted in different colors corresponding to the different enriched Pathways in Gene ontology, KEGG and Reactome databases. **(C)** q-PCR validation of representative transcripts (*OASL, IFI6*) of 13-genes signature in OCI-AML3, *MDS-L*, and F36P cells after CM-272 treatment. Bar graph represents fold change expression [mean ± standard deviation (s.d)] of 3 independent experiments performed in technical triplicate. Welch’s *t*-test was performed for *p-* value statistical significance. **(D)** Heatmap depicting ISMARA TF motif analysis (based on *z* score of top predictor motif activities, TF-gene Pearson correlation, and average gene target expression change). **(E)** BubbleGUM gene set enrichment analysis (GSEA) map for inflammatory pathways of OCI-AML3 RNA-seq established from MsigDB database. For each gene set, origin of the data set and number of upregulated (red, CM272 treated) and downregulated (blue, DMSO control treated) genes are indicated. The panel summarizes the normalized enrichment score (NES) and FDR parameters. **(F)** ERV retroelements expression in OCI-AML3 post-CM272 treatment (RNA seq, three biological replicates for each condition). Data are presented as Log2 Fold-change ± s.d. **(G)** Cytokine release profile in the supernatant of OCI-AML3 post-CM272 treatment (200nM for 48h) and determined by Luminex assays. Data are presented as mean of protein detection ± s.d. from 3 independent experiments. **(H)** Distribution of DNA methylation changes in relation to CpG islands (CGI), including shores, shelves, and open sea regions (inter-CGI) for differentially hyper- or hypomethylated CpG sites in OCI-AML3 post-CM-272 treatment compared with the DMSO control. Random bar shows the distribution of randomly selected (n=10.000) CpG sites from EPIC Methylation Array. (**I)** Enrichment analysis of differentially CpG sites (DMCpGs) located in different genomic regions, annotated by 15 chromHMM states. Color scale refers to log2 fold change and circle size refers to -Log10(FDR) *q*-value significance. **(J)** Gene Ontology term enrichment pathways of hypomethylated DMCpGs using GREAT analysis tool. **(K)** Bubble plot representation of HOMER transcription factor (TF) motif enrichment analysis of differentially hypo- and hypermethylated DMCpGs (left and right panel, respectively). Color range depicts different transcription factor families and circle size refers to -log10(*p*-value) significance.

To characterize the NPM1c AML transcriptional regulation of these altered gene programs in more detail, we performed an inferred motif-centric transcription factor activity analysis (ISMARA) ^46^ on RNA-seq data of OCI-AML3. In response to CM-272, top transcriptional regulators were identified among three categories of transcription factor (TF) motifs with increased activity (Figure 4D): common myeloid regulators (SPI1, ATF3), mature myelo-monocytic modulators (RUNX1, EGR1, JUNB) and type I IFN response instigators (IRF3, IRF7, IRF9) ^47–49^. Inversely, TF motifs for oncogenes (MYC, HOXA9, MEIS1) were found to be less active in response to CM-272 **(Figure 4D)**, thereby reversing the aberrant transcriptional program of known oncogenic drivers and launching a viral mimicry response. Furthermore, targeted GSEA analysis of OCI-AML3 RNA-seq data confirmed up-regulation of pro-inflammatory and IFN pathways linked to antiviral defense in response to CM-272 while decreasing the activation of E2F-dependent cell cycle signaling **(Figure 4E and supplementary Figure S5D-E)**. As recent studies showed that Azacytidine treatment in MDS/AML can trigger viral mimicry response by re-expression of retrotransposons ^50,51^, we analyzed the RNA-seq data of OCI-AML3 for level changes in double-stranded (ds) RNA transcribed by long terminal-repeat sequences (LTR). In response to CM-272, 89 LTRs were found to be substantially re-expressed, been mostly of endogenous retroviral (ERV) origin (Figure 4F), including LTR16A1, MLT2A2, and LTR12C, which belong to the ERV1 superfamily **(supplementary Figure S5F)** ^50,52,53^. Another consequence of viral mimicry response involves the activation of IFN and pro-inflammatory signals ^54^, which was confirmed by detection of elevated levels of IL-1α, IL-8, IFN-γ and TNF-α in the supernatant of OCI-AML3 following CM-272-treatment **(Figure 4G)**.

Since most of the retrotransposons are tightly regulated by DNA methylation, we generated a whole epigenome profile of OCI-AML3 to determine the extent of chromatin remodeling in response to CM-272. Similar to known decitabine effects, global hypomethylation was observed after 96h of CM-272 treatment along with occurrence of specific localized hypermethylation **(supplementary Figure S5F)**, a phenomenon also observed during monocytic differentiation or after HMA treatment of DNMT3A^mut^ cells ^55,56^. From a total of 9,708 differentially methylated CpGs (DMCpGs), 8,380 sites (86%) were hypermethylated and 1,328 sites (14%) were hypomethylated **(supplementary Figure S5G)**. We next analyzed the distribution of DMCpGs in relation to the CpG islands known to be enriched in the silenced genome. In response to CM-272, both hypomethylated and hypermethylated DMCpGs were located primarily in open sea areas (interCGI) but not in CpG islands **(Figure 4H)**. Using a 15-state chromatin model for K562 AML cells ^57^, we showed that hypomethylated DMCpGs were significantly enriched in active states (associated with expressed genes) which consist mostly of active transcription start site (TSS) proximal promoter states (TssAFlnk) and enhancer regions (EnhG, Enh)., By contrast, hypermethylated DMCpGs were significantly enriched in quiescent regions of inactive states (associated with silenced genes) **(Figure 4I)**. Additionally, Gene Ontology (GO) analysis of genes associated with differentially hypomethylated sites were enriched in biological processes related to regulation of monocyte differentiation and defense immune response **(Figure 4J)**. This further supports the idea that CM-272 treatment enhances myeloid differentiation. To investigate if these mechanism-associated loci are mutually regulated, we performed a search for enriched transcription factor (TF)-binding sites in these DMCpG regions using the HOMER algorithm ^58^. In the hypomethylated regions, we observed a significant over-representation of binding sites for the bZIP and ETS family that are linked to myeloid lineage (PU-1, ERG) and mono-macrophage differentiation (MAFB, BACH-1) as well as transcriptional activators (AP-1, ATF, JUNB). In the hypermethylated regions on the other hand, we observed weak enrichment of E2F, IRF and Runt family members of TFs related to cell cycle regulation and dendritic cell differentiation from the monocytic lineage **(Figure 4K)**. These results suggest that key transcription factors involved in the upregulation of myelo-monocytic genes are aberrantly silenced by DNA hypermethylation in NPM1c AML.

Overall, our data revealed that remodeling of the epigenome by the inhibition of DNMT/G9 complex in AML cells predominantly involves very specific alterations of the methylation status that play an important role in the myeloid differentiation process toward monocytic lineage but does not alter larger chromatin regions.

### In vivo efficacy of CM-272 monotherapy against NPM1c AML cells

To assess the in vivo efficacy of CM-272 against NPM1c AML, we utilize an OCI-AML3 xenograft model. To this end, immunodeficient NBSGW mice were transplanted with cells carrying Luciferase/GFP tags for bioluminescence imaging and flow cytometry analysis (OCI-AML3-luc-GFP). After successful engraftment, mice were treated with vehicle or CM-272 **(Figure 5A)** ^32,33^. Compared to vehicle, CM-272 monotherapy for 4 weeks significantly reduced OCI-AML3-luc-GFP tumor burden in NBSGW mice **(Figure 5B)** across different tissues including the liver **(Figure 5C)**, known to be infiltrated in aggressive refractory/relapse AML settings ^59,60^. Leukemia reduction was mostly driven by the cytotoxic effect on engrafted OCI-AML3-luc-GFP highlighted by the massive depletion of tumor cells in infiltrated liver tissues as shown with flow cytometry **(Figure 5D-E)** and pathological analysis **(Figure 5F-G)**.

**Figure 5.**
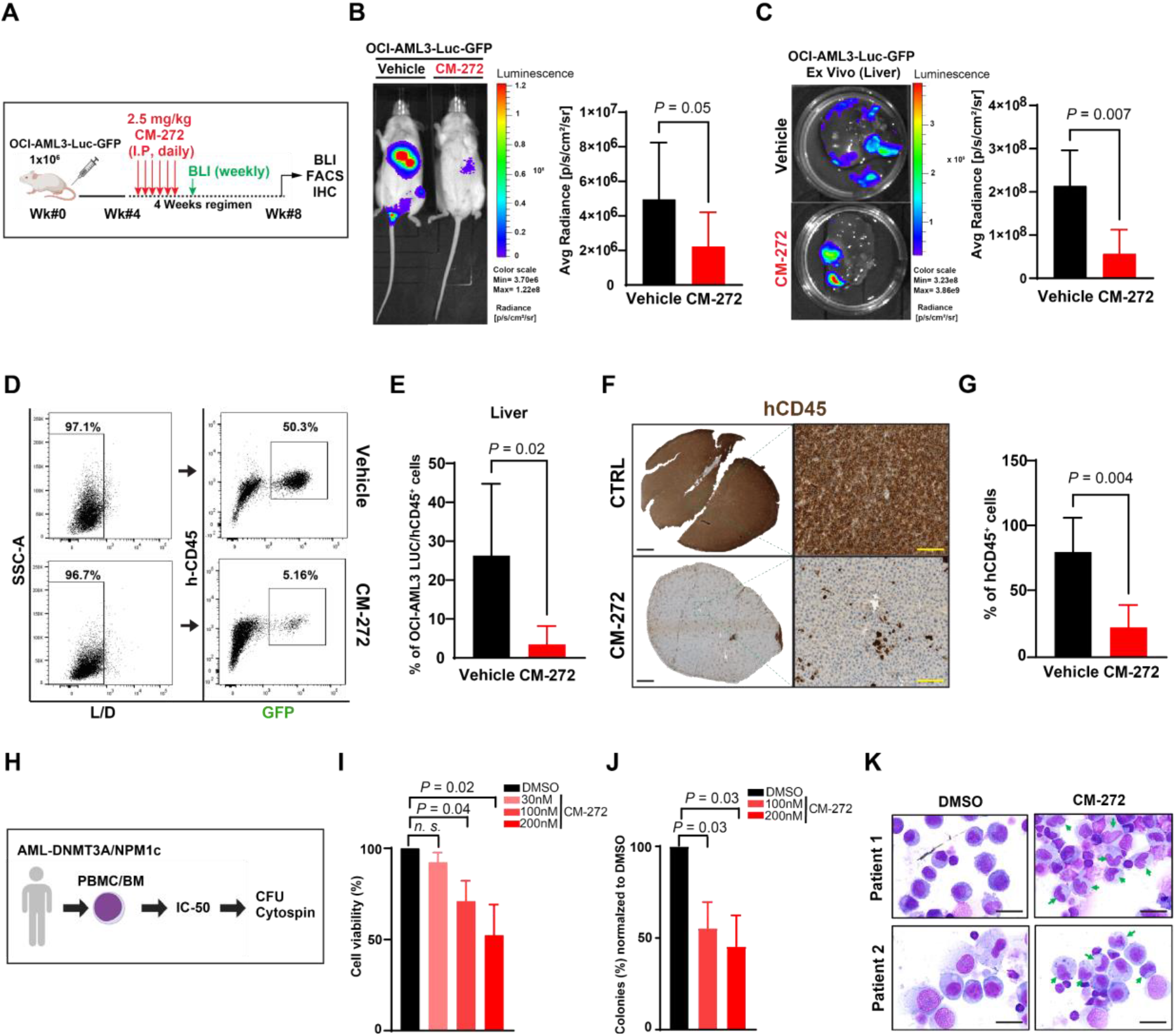
Anti-tumor growth efficacy of DNMT/G9a dual inhibitor in DNMT3A^mut^ NPM1^c^ AML mouse model and patient-derived samples. **(A)** Experimental design of *in vivo* administration of CM272 in AML xenograft mouse model. NSGW41 mice were injected intravenously with OCI-AML3-Luci-GFP cells (1.0×10^6^). Leukemia engraftment was confirmed 1 week later through a non-invasive *in vivo* bioluminescence imaging (BLI) system following injection with a D-luciferin (4 mg/mouse) substrate. Mice were dosed daily with an intravenous vehicle or an intravenous active form of CM-272 (2,5 mg/kg, 5 days/week) up to 4 weeks. **(B)** Representative full body BLI data for each condition at week 8. Bar graph represent Luciferase intensity (mean ± s.d) at week 8 for n = 8-9 mice per condition. Unpaired t test was performed for *p-*value statistical significance. **(C)** Representative Liver BLI data for each condition at week 8. Bar graph represent Luciferase intensity (mean ± s.d) at week 8 for n = 5 liver per condition. Unpaired t test was performed for *p-*value statistical significance. **(D)** Representative FACS histogram of livers isolated from xenografted recipient mice with OCI-AML3-Luci-GFP at week 8. Squares and arrows indicate gating strategy to identify CD45^+^-GFP^+^ xenografted cells in each condition. **(E)** Bar graph represent percentage of xenografted OCI-AML3-Luci-GFP in NSGW14 mice (mean ± s.d) at week 8 for n = 5-7 mice per condition. Unpaired t test was performed for *p-*value statistical significance. **(F)** Representative liver sections of human CD45 engraftment in recipient mice with OCI-AML3-Luci-GFP at week 8. huCD45 positive signal is assessed by Immunohistochemistry staining with chemiluminescence (HRP). Micrographs were taken at x10 and x60 amplification. **(G)** Bar graph represent percentage of xenografted human CD45 engraftment in recipient mice (mean ± s.d) at week 8 for n = 4-6 mice per condition. Unpaired t test was performed for *p-*value statistical significance. **(H)** Experimental design of CM-272 treatment on primary bone marrow and peripheral samples from *de novo* DNMT3A^mut^ NPM1^c^ (+/-FLT3-ITD) patients. AML blasts was treated with CM-272 (30, 100 and 200nM) and DMSO control for 72h **(I)** Cytotoxicity of CM-272 was measured on five independent samples of DNMT3A^mut^ NPM1^c^ AML. Cell viability was assessed with DAPI staining by flow cytometry and performed in triplicates. Data are presented as percentage mean of viable cells ± s.d. **(J)** CFU assays were carried out in technical duplicates and scored after 8 days incubation in methylcellulose. Data are presented as colonies percentage normalized to DMSO control ± s.d. **(I-J)** One-way ANOVA was performed for *p-*value statistical significance. **(K)** Giemsa-stained cytospins showing Leukemic cells from two DNMT3A^mut^ NPM1^c^ AML patients and 7 days after single treatment with CM-272 (200nM) in comparison to DMSO control. Micrographs were taken at x100 amplification arrows indicate morphological changes.

Finally, we tested the efficacy of CM-272 in primary samples from DNMT3A^mut^ NPM1c AML patients **(Figure 5H)**. The majority (four out of five) of these samples also carry FLT3-ITD mutations **(Supplementary Table S2)**, a mutational recurrence and a common feature that generally confers a relatively poor prognosis 2. Overall viability **(Figure 5H and supplementary Figure S6A)** and colony number was significantly decreased **(Figure 5J)** along with increased apoptosis **(supplementary Figure S6B)** which is dependent on escalating concentrations of CM-272. More strikingly and similar to OCI-AML3 **(Figure 2I)**, morphological signs of myelo-monocytic differentiation were also detected in primary AML cells post-CM-272 treatment **(Figure 5K)**. Together, these data demonstrate that, regardless of the nature of the cooperating mutation, all tested human primary NPM1c leukemias were highly sensitive to growth inhibition by CM-272.

In conclusion, we propose that dual targeting of DNMT/G9a epigenetic activities represents a so far unrecognized treatment option that should be further tested in vivo to improve the clinical management of DNMT3A/NPM1-mutated AML with or without FLT3-ITD.

## Discussion

AML is a molecularly complex disease characterized by heterogeneous genetic tumor profiles and involving numerous pathogenic mechanisms and pathways. Integration of molecular data across multiple clinical cohorts has advanced our current understanding of AML pathogenesis and genetic classification toward a better patient risk stratification ^61^. In recent years, the classification of patients using DNA methylation patterns allowed to identify distinct AML subgroups, such as those harboring NPM1 or IDH1/2 mutations ^62–64^, which led to widespread optimism in the development of epigenetic therapies as an additional or complementary arm to current cytotoxic chemotherapy regimens ^65^. Nevertheless, there are still unmet clinical needs to overcome resistance and low responsiveness toward these epigenetic therapies to yield durable remissions and extend patient survival.

In this study, we discovered and fully characterized a specific epigenetic vulnerability of AML with DNMT3A/NPM1 mutations **(Figure 1 and 2)**. This specificity appears to originate from the particular methylation patterns affecting mostly promoter CpG islands that have been highly associated with mutated DNMT3A in clonal hematopoiesis and overlapping in AML with the co-occurrence of NPM1c ^66,67^. Accordingly, we demonstrated a significant association of these promoter CpG regions and the regulation of myelo-monocytic differentiation in response to the dual DNMT/G9 inhibitor CM-272 **(Figure 3)**. We showed that this control mechanism is driven by a dynamic epigenetic rearrangement that is not restricted to loss of methylation, as previously reported for HMA therapy, but commonly involves gain of methylation ^56,68^.

Our study further reveals that remodeling of the epigenome during CM-272-induced differentiation predominantly involves local tweaking of the methylation status rather than global changes across larger hypomethylated regions **(Figure 4)**. Most of these hypomethylated regions were enriched for multiple binding sites of TFs of the ETS (PU.1, ERG, MAFB) and bZIP motif family (AP-1, ATF3, JUNB) and containing mainly AP-1 TFs that are known to be key players in monocytic differentiation and active transcription sites ^69,70^. Moreover, chromatin changes associated with dual inhibition of DNMT/G9a are enriched in genomic regions associated with enhancers. This is reminiscent of G9a-dependent de novo CpG methylation of pluripotent genes, which require the active recruitment of DNMT3A for transcriptional repression toward control of embryonic development ^71,72^. Surprisingly and despite the relocation of NPM1 under CM-272 treatment, we didn’t observe major changes in HOX genes expression, which is consistent with recent studies using NPM1-Crispr genetic deletion and Menin inhibitors ^73,74^. This could be partially explained by previous reports, showing that G9a controls the activity of HOXA9 via direct physical interaction in AML, which can be deregulated by a G9a-specific inhibitor, thereby inducing differentiation ^20,75^. In parallel, the observed nuclear relocation of cytoplasmic mutant NPM1, which also controls aberrant HOX gene expression, can be attributed to a decrease

## Materials and Methods

### Small molecule inhibitors

Hypomethylating agent (Decitabine, DAC), DNMT inhibitor (SGI-1027) and dual DNMT/G9a inhibitor (CM-272) were purchased from in gene expression of XPO-1 **(Figure 3I)**, since XPO-1 acts as a nuclear exporter for mutant NPM1 ^76^. Furthermore, the NPM1c relocation can be explained by a rearrangement of chromatin accessibility via 3D genome reorganization within the nuclear membrane, known to be abundant with H3K9me2 epigenetic marks that control key regulatory regions for HOX genes and myeloid differentiation ^77,78^.

Previous reports demonstrated that both DNA methylation and H3K9 methylation are involved in the formation and maintenance of constitutive heterochromatin and the silencing of retrotransposons ^79,80^. Our data indicate that treatment with CM-272 results in de-repression of specific ERVs from the HERV-L family **(Supp Figure S5F)** that are known to be under the dual control of DNA and H3K9me2 methylation ^81,82^. This triggers the viral sensing machinery and interferon-mediated cell death as shown with previous studies using genetic deletion of other members of the HMT family such as SETDB1 and SUV39H1 ^83,84^. In addition, previous reports showed that DNMT inhibition or genetic HMT deletion alone showed incomplete ERV re-expression ^50,85^, thereby reinforcing the concept of dual DNMT/G9a targeting to promote viral mimicry response toward activation of immunogenicity.

Being of considerable translational importance, we demonstrated that dual DNMT/G9a inhibition acts on differentiation induction and interferon-mediated cell death leading to significantly reduced tumor growth in clinical AML samples **(Figure 5)**. We believe that our results are robust enough to warrant further optimization in future studies with potential combination with other currently tested therapies on NPM1c AMLs such as Venetoclax or Menin inhibitors in future studies. Overall, our approach to target epigenetic vulnerabilities in specific AML subgroups opens a new avenue for refined therapeutic strategies with a new class of chromatin modulators. Indeed, successful combinations of epigenetic drugs and immune checkpoint inhibitors in animal models and early clinical trials suggest that these novel drugs have immunomodulatory properties. So far, the effects of these epigenetic inhibitors appear to be mediated, at least in part, through the reactivation of non-coding elements of the genome that remain to be completely understood before incorporating them into AML clinical regimens.

MedChemExpress LLC. For *in vitro* studies, DAC, SGI-1027 and CM-272 were dissolved in DMSO, aliquoted and stored at −80°C. For *in vivo* applications, CM-272 was dissolved in DMSO and added freshly to PBS prior to intraperitoneal injections.

### Cell lines and culture conditions

OCI-AML3, OCI-AML2, MDS-L, F-36P and all other human AML cell lines used in this study were purchased from German Collection of Microorganisms and Cell Cultures GmbH (DSMZ), tested at regular periods for mycoplasma and authenticated by Single Nucleotide Polymorphism (SNP)-profiling (Multiplexion, Heidelberg, Germany), as described previously ^86^. All AML cell lines were maintained in standard culture media conditions, as described previously ^87,88^. The murine retrovirally transformed Hoxa9-Meis1 bone marrow cells (kindly donated by Michael Kühn lab, UMC Mainz) and their culture conditions have been described elsewhere ^40^.

### Plasmid Generation, Viral Packaging, and Creation of OCI-AML3-Luc-GFP Line

The firefly luciferase (Fluc) expression plasmid pLenti-EF1a-Fluc-P2A-EGFP-T2A-PAC was generated by replacing the Gaussia luciferase (GLuc) of the pLenti-PAC-T2A-GLuc plasmid (kindly provided by Preet Chaudhary, University of Southern California, CA, USA) with the firefly luciferase hluc+ (Promega, Waltham, MA, USA). PCR products encoding FLUC, EGFP and PAC were assembled in frame using Gibson Assembly (NEB, MA, USA), separated by the ribosomal skipping element P2A and T2A, respectively, in the order Fluc-P2A-EGFP-T2A-PAC. Sequencing was performed using the services of Eurofins (Luxembourg, Luxembourg). Sequence analysis was performed using Geneious software (Biomatters, New Zealand).

Plasmid constructs for the production of lentivirus were transfected with packaging plasmids psPAX2 and pMD2.G into HEK-293FT cells utilizing TransduceIT™ Transfection Reagent (Mirus Bio LLC, WI, USA) according to the manufacturer’s instructions. Media was changed the following day. Viral supernatant was collected 72 hours post transfection and filtered through a 0.45 µm PES membrane. OCI-AML3 cells were spinofected with virus particles (60 min at 2000 rpm) in the presence of 5µg/ml polybrene (Sigma Aldrich). The following day, the viral supernatant was removed by centrifugation and cells were transduced with fresh viral supernatant for an additional 24 hours. Positive OCI-AML3-Luc-GFP cells were then selected with 1µg/ml puromycin (Sigma Aldrich) and stable expressing cells were maintained in long-term culture under selection prior to their utilization in therapeutic in vivo mouse studies.

### Primary AML biopsies and blood samples

All primary samples used in this study (liquid and paraffin sections) were obtained under institutional review board–approved protocols and in accordance with the Declaration of Helsinki. Formaldehyde-fixed paraffin-embedded (FFPE) precut sections of human bone marrow biopsies from *NPM1*^*mut*^ *FLT3*^*ITD*^ AML samples (n=15) were provided by the Tissue bank of the University Hospital of the Technical University of Munich (MTBIO). Healthy FFPE bone marrow biopsies (n=10) were provided by the University Hospitals in Dresden and Leipzig as part of the BoHemE study (NCT02867085). Cryopreserved primary peripheral blood or bone marrow blasts from *DNMT3A*^*mut*^ *NPM1*^*mut*^ *FLT3*^*ITD*^ AML samples (n=5) were obtained from patients treated at the University Medical Center, Mainz. Additional information on AML patient samples used in IHC and in vitro assays is available in **Supplementary Table S1 and S2**.

Umbilical cord blood (UCB) samples were obtained from scheduled cesarean section of normal full-term deliveries with informed consent from the Gynecology Department, University Medical Center Mainz. Mononuclear cells (MNC) were isolated by Ficoll-Paque (Millipore Sigma) density gradient centrifugation as reported elsewhere ^89^. After erythrocyte removal (ACK lysis, Gibco Thermo Fisher), CD34^+^ cells in the MNC-UCB fraction were isolated using CD34 MicroBead Kit UltraPure (130-100-453, Miltenyi Biotec) via LS Columns and MidiMACS Separator (Miltenyi Biotec). CD34^+^ cells were counted, assessed for depletion quality by flow cytometry and cryopreserved in liquid nitrogen for other assays.

### Drug sensitivity assays and cell viability assessment

Cells were plated in 96-well plates at a density of 5×10^3^ cells per well and exposed to increasing drug concentrations. After 24 or 72 h, cell viability was determined with a CellTiter-Glo Luminescent Cell Viability Assay according to the manufacturer’s description (G7570, Promega) on a FLUOstar Omega microplate reader (BMG Labtech GmbH). Absolute absorbance values were normalized to DMSO control treatment and IC_50_ values were computed in GraphPad Prism version 9.2.0 with nonlinear fit of log(inhibitor) vs. response - Variable slope (four parameters). All experiments were tested in technical triplicates and each experiment was performed at least three times.

Primary cells were seeded in 12-well plate at a density of 1×10^6^ cells per well in 2 ml StemSpan™SFEM II media (STEMCELL Technologies) with growth factors cocktail (100 ng/ml SCF, 50 ng/ml FLT3-L, 20 ng/ml IL-3 and 20 ng/ml G-CSF; all purchased from Peprotech) supplemented with 1% penicillin–streptomycin (ThermoFisher Life Technologies) and 1 μM of StemRegenin 1 (Selleckchem LLC). On the same day, the cells were treated with 30 nM, 100 nM, 300 nM, 500 nM, or 1000 nM CM-272 or vehicle (DMSO) for 72 h. Apoptotic and viability effects of CM-272 on treated primary AML cells was investigated by the Caspase-Glo 3/7 assay (G8981, Promega) following the manufacturer’s instructions and measured in the FLUOstar Omega microplate reader. To analyze only the Caspase 3/7 activity of viable cells, the cell viability assay CellTiter-Glo was performed simultaneously. Luminescence signals of the Caspase 3/7 assay were normalized to the luminescence signal of CellTiter-Glo. Drug-treated samples were normalized to the DMSO control to determine the percentage of viable cells affected by CM-272. For each primary sample, all experiments were performed in technical triplicates.

### Flow cytometry assays and cell surface marker analysis

To characterize myeloid differentiation status, OCI-AML3 cells (2×10^6^ cells/well) seeded in 2 ml of culture medium were exposed to 200 nM of CM272 for various timespans (24, 48, 72, 96h and 120h). Cells were then harvested, washed and stained with anti-CD11b-APC (clone ICRF44, 17-0118-42, ThermoFisher Life Technologies) and anti-CD14-PE (clone 63D3, 367130, BioLegend). Unstained cells and respective PE and APC Isotype controls were used as controls. Next, cells were washed twice (300 g, 4 min, 4°C), resuspended and incubated in FACS buffer (1×PBS, 2 mM EDTA and 0,1% BSA) containing 0.1 µg/ml of DAPI 10 min before analysis. The effects of CM-272 on apoptosis (cleaved PARP), DNA damage (γ-H2AX) and cell cycle (BrdU) after 24 and 72 hours of treatment were determined by flow cytometry using the BD Pharmingen Apoptosis, DNA Damage, and Cell Proliferation Kit (562253, BD Biosciences) according to the manufacturer’s instructions. All flow cytometry analyses were performed on BD FACSCanto II (BD Biosciences) and subsequent analyses were done using FlowJo v10.8 Software (BD Biosciences). All experiments were performed in technical triplicates and each experiment was performed at least three times.

### Colony-forming unit (CFU) assay

In order to quantify the colony formation capacity of cells under different CM-272 concentrations, 500 cells/1ml (AML cell lines and CD34^+^ UCB) or 5000 cells/1ml (primary AML samples) were seeded in MethoCult™ H4034 Optimum or Methocult H4230 (Stemcell technologies) supplemented with 10% FBS, 1% penicillin-streptomycin, cytokines (50 ng/ml SCF, 10 ng/ml IL-3, 10 ng/ml GM-CSF, 3 U/ml EPO) and the indicated CM-272 concentrations into six-well plates (SmartDish™ 27370, Stemcell technologies). Cells were cultured for 8-10 days at 37 °C and 5% CO2 and colonies (defined as a collection of at least 10 cells) were quantified and pictured using an EVOS M5000 imaging system (ThermoFisher) at 20x magnification. All experiments were performed in technical triplicates and each experiment was performed at least three times.

### Cytokine measurement assay

Human Custom ProcartaPlex Luminex panel assay (Bio-Techne) was used to measure the levels (pg/ml) of 16 analytes including TNF-α, IL-6, IL-8, IL-1α, and IFN-γ in the supernatants of OCI-AML3 following 48h CM-272 treatment. All experiments were performed in technical triplicates and each experiment was performed at least three times.

### Western-blot analysis

Cells treated with drug(s) or vehicle were lysed in RIPA-buffer (8990, ThermoFisher) supplemented with protease and phosphatase inhibitor cocktail (11836170001, Roche). For the subcellular localization of the mutated NPM1c within OCI-AML3 after DMSO or CM-272 treatment (72 h), cell fractionation was performed before cell lysis using cell fractionation kit (9038, Cell Signaling Technology) and following the manufacturer’s instructions. In all cases, the protein concentration was determined with a Pierce BCA Protein Assay Kit (23227, ThermoFisher). Equal amounts of protein were used for gel electrophoresis and blotted on a nitrocellulose PVDF membranes (Mini-PROTEAN system, Bio-Rad). Membranes were cut to allow independent incubation with antibodies for 60-90 min or overnight. Following antibodies were used: anti-DNMT1 (1:1000, 5032), anti-DNMT3A (1:1000, 3598), anti-DNMT3B (1:1000, 67259), anti-G9a (1:1000, 68851, all purchased from Cell Signaling Technology), anti-Meis1 (1:500, MA527191, ThermoFisher), anti-PBX3 (1:500, 12571-1-AP, Proteintech), anti-cMYC (1:2000, 10828-1-AP, Proteintech), anti-H3K9me2 (1:1000, ab1220, Abcam), anti-H3 (1:1000, ab1791, Abcam) and anti-GAPDH (1:10000, AM4300, ThermoFisher). In addition, anti-mutant NPM1c (1:500, PA146356, ThermoFisher), anti-Vinculin (1:1000, 13901, Cell Signaling Technology) and anti-Lamin B1 (1:1000, 12987-1-AP, Proteintech) were used for subcellular fraction analysis. After washing steps, membranes were incubated for 1h with HRP-conjugated goat secondary antibodies against mouse or rabbit immunoglobulin (1:3000, 170-6516, 170-6515, Bio-Rad). Proteins were visualized on an iBright™ CL1000 Imaging System (ThermoFisher) using SuperSignal™ West Pico Plus Chemiluminescent Substrate (34578, ThermoFisher) following manufacturer’s instructions. Densitometry analysis was performed using iBright Analysis Software (ThermoFisher). All experiments were performed at least twice. All images have been cropped for improved clarity and conciseness.

### In vivo study of CM-272 in AML xenograft mouse model

NOD.Cg-Kit^W-41J^ Tyr^+^ Prkdc^scid^ Il2rg^tm1Wjl^/ThomJ (NBSGW; Jax#026622) were injected in the lateral tail vein with 1×10^6^ OCI-AML3-Luc-GFP cells. Mice were imaged utilizing an IVIS Lumina II in vivo imaging system (PerkinElmer) to document engraftment before treatment was initiated. Briefly, mice were anesthetized and imaged noninvasively after intraperitoneal injection of D-Luciferin (15 mg/mouse) (AppliChem, Darmstadt). Five weeks thereafter, xenografted NBSGW mice were randomized into groups with equivalent mean bioluminescent intensity and subsequently subjected to different treatments: Group 1 (n=9, CM-272, 2,5 mg/kg in PBS, daily x 6 days, by intraperitoneal injection) and Group 2 (n=8, Vehicle, PBS alone, same regimen). All mice were treated up to 4-5 weeks and imaged weekly by bioluminescent imaging to document treatment efficacy and/or disease progression. Total bioluminescent flux was recorded as photons/second/cm^2^/steradian and analyzed with Living Image Software 4.0 (PerkinElmer). Mice that became moribund were euthanized according to the approved IACUC protocol and as final end-point all experiments were terminated 8 weeks post-transplant. Liver, spleen and bones were harvested and individually analyzed for bioluminescent intensity. Thereafter, tissues were processed for engraftment assessment by FACS, with gating for GFP^+^ human CD45^+^ cells (clone 2D1, 17-9459-42, ThermoFisher). Immunohistochemistry analysis on liver sections was performed as described in the section below. NBSGW mice were housed and treated at the animal facility of the Institute for Tumor Biology and Experimental Therapy (Frankfurt am Main, Germany) in accordance with regulatory guidelines. All animal experiments were reviewed and approved by the Regierungspräsidium Darmstadt, Germany under protocol number F123/1057.

### Immunohistochemistry

3-5 µm sections of human FFPE-bone marrow AML biopsies were prepared on which immunohistochemistry (IHC) was performed either based on chromogenic-labeling (DAB) or immunofluorescent-labeling (IF). Deparaffinization, rehydration and antigen retrieval were performed by heating sections in Trilogy™ buffer (Cell Marque, Millipore Sigma) followed by peroxidase blocking for IHC-DAB (0.3% H2O2 in methanol). For all IHC applications, permeabilization was done with 0.2% Triton X-100 (Millipore Sigma) in PBS. Next, sections were blocked with 5% donkey or goat serum (Jackson ImmunoResearch Europe Ltd) in PBS and incubated overnight with primary antibody as follow: rabbit anti-H3K9me2 (1:500, Ab176882, Abcam) for IHC-DAB analysis; mouse anti-H3K9me2 (1:500, ab1220, Abcam) and rabbit anti-G9a (1:200, HPA050550, Millipore Sigma) for IHC-IF. Slides were then rinsed with PBS and chromogenic staining was performed using Rabbit VisUCyte HRP Polymer-DAB Cell & Tissue Staining Kit (VCTS003, R&D Systems) following manufacturer’s instructions and counterstained with hematoxylin (Millipore Sigma). For IHC-IF, slides were incubated with Alexa Fluor™ 647 or 568-conjugated donkey secondary antibodies against mouse or rabbit IgGs (1:1000, Invitrogen, ThermoFisher) and after washing, slides were mounted using Fluoroshield solution (ab104135, Abcam). For analysis of liver tissues from OCI-AML3-Luc xenografted NSGW41 mice, IHC-DAB for human CD45 expression (M0701, DAKO, Leica) was performed as previously described ^60^. Thereafter, DAB and IF-stained slides were scanned up to original magnification ×40 on Scanscope CS2 (Aperio, Leica) and Opera Phenix High-Content Imaging System (PerkinElmer) respectively. Fluorescence quantification was performed with Harmony high-content analysis software (PerkinElmer), while DAB quantification was analyzed using QuPath software as described previously ^90,91^.

### Immunofluorescence and morphological analysis

For subcellular localization of NPM1-wt and mutant, 1×10^6^ OCI-AML3 cells were resuspended after 48 h CM-272 treatment in 150 μl PBS and applied onto coverslips (precoated with 0.01% poly-l-lysine; Millipore Sigma) in a humidified chamber for 10 min. Cells were fixed with 4% paraformaldehyde and permeabilized using PBS with 0.2% Triton X-100. Slides were incubated with Image iT-Signal Enhancer (I36933, ThermoFisher) at RT for 30 min prior to immunostaining. After washing, coverslips were incubated with rabbit anti-mutant NPM1/c-terminal (1:100, ABIN1737584, antibodies-online) and mouse anti-wild-type NPM1/B23 (1:50, sc-70392, Santa Cruz) diluted in PBS-0.2% Tween-20 (Sigma) for 2 h at RT. After washing steps, coverslips were incubated with Alexa Fluor™ 488 or 647-conjugated goat secondary antibodies against mouse or rabbit IgGs (1:1000, Invitrogen, ThermoFisher) and then mounted using Fluoroshield solution (ab104135, Abcam). Cells were pictured using an EVOS M5000 imaging system (ThermoFisher) at 100x magnification, then fluorescence was quantified with ImageJ analysis software (National Institutes of Health). Each experiment was performed at least twice.

For morphological analysis post-CM-272 treatment, 1×10^5^ cells from CFU or in vitro cultures were harvested in 150 μL PBS. Cells were centrifuged using a Cytospin 4 (ThermoFisher) onto coated cytoslides (5991056, ThermoFisher) at 500 rpm for 5 min. The cells were fixed and stained with May-Grünwald followed by a 4% Giemsa staining (both from Millipore Sigma). Cellular/nuclear morphology was assessed by light microscopy and pictured using an EVOS M5000 imaging system (ThermoFisher) at 60x magnification. Each experiment was performed at least twice.

### RNA purification, cDNA synthesis and qPCR

Total RNA was isolated using the PureLink RNA Mini Kit (Ambion, ThermoFisher) for downstream experiments including quantitative real-time polymerase chain reaction (qPCR) and RNA sequencing (RNA-Seq). Samples for RNA-Seq were treated with an extra on-column DNase I digestion step. cDNA synthesis was performed using qScript cDNA SuperMix Synthesis Kit (Quantabio) and qPCR were carried out using PowerUp™ SYBR™ Green MasterMix (ThermoFisher) for human sequences or TaqMan Fast Advanced MasterMix (ThermoFisher) for murine sequences, and run on QuantStudio3 Thermocycler (ThermoFisher). Relative gene expression was determined by the 2-ΔΔCT method and normalized to the internal control, β-2 microglobulin or GAPDH. Primer sequences are available on request. All experiments were performed in technical triplicates and each experiment was performed at least three times.

### RNA sequencing (RNA-seq) and gene expression analysis

RNA was quantified and checked for integrity using Qubit Fluorometer (Thermofisher) and an Agilent 2100 Bioanalyzer (Agilent). Libraries were prepared using TruSeq stranded mRNA Library Prep kits (Illumina) following manufacturers’ recommendations. The libraries were sequenced on an Illumina NextSeq500 machine in single end mode with 75 bp read length yielding an average of 25 million raw reads per sample. RNA-seq alignment and specialized bioinformatics analyses were performed with the support of the Bioinformatics Core Facility (Institute of Molecular Biology, Mainz). Assessment of raw data quality, rRNA content and PCR duplication rates was done using FastQC^92^, FastQ Screen^93^ and the R package dupRadar v.1.16.0^94^. For gene expression analysis, reads were mapped to the human reference genome Hg38 (GRCh38, gencode release 35) using STAR^95^ and summarized at gene level using Subread featureCounts v1.6^96^. Pairwise differential expression analysis between the experimental groups of CM272 treated cells vs. DMSO treated controls (3 replicates each) for all 3 cell lines (OCI-AML3, F-36P and MDS-L) was performed with the R-package DESeq2 v.1.26.0^97^. The analysis was performed in paired sample design, which includes the sample (replicate) information as a term in the DESeq2 design formula. Due to high similarities in gene expression profile with the MDSL_DMSO control group, the sample pair MDSL_CM272_2 (replicate 2) and MDSL_DMSO_2 (replicate 2) has been removed from downstream differential expression analysis, leaving 2 replicates per condition for this group comparison. Genes were tested for an absolute fold change threshold of >2 specified in the DESeq2 statistical testing model. P-values were calculated by a two-tailed Wald test for testing if the observed fold-changes are greater in absolute value than the given threshold, which will result in more conservative estimates compared to post hoc filtering. Genes are considered as significantly differentially expressed if the obtained p-value for this test is below 0.01 after adjustment for false discovery rate (FDR, q-value < 0.05). Sets of significant up-regulated and down-regulated genes are analyzed for over-represented gene ontology terms (GO) and biological pathways as implemented in the R-packages clusterProfiler v.3.14.0^98^ and ReactomePA v.1.30.0^99^ applying a significance threshold of p < 0.01 after adjustment for FDR.

### Downstream analyses of RNA-seq (GSEA, TE and ISMARA analyses)

Gene set enrichment analysis (GSEA) comparing two groups (DMSO vs CM-272) was performed using GSEA software v4.1.0 and public gene sets downloaded from MSigDB (https://www.gsea-msigdb.org). High-throughput GSEA were performed using the BubbleMap module of BubbleGUM software^100^. BubbleMap analysis was performed with 1000 gene set–based permutations and with “Signal2noise” as a metric for ranking the genes. The results are displayed as a bubble map, where each bubble is a GSEA result and summarizes the information from the corresponding enrichment plot. Red bubbles indicate enrichment in CM-272 treatment and blue bubbles enrichment in DMSO-treatment. Strength of enrichment is represented by normalized enrichment score (NES). Significance of enrichment is represented by the false discovery rate (FDR, q-value < 0.05) derived by computing the multiple testing–adjusted permutation-based P value using the Benjamini-Yekutieli correction.

Quantification of both gene and transposable element (TE) transcript abundances from RNA-Seq experiments was performed with TEtranscripts software package^101^ and utilizing both uniquely and ambiguously mapped short read sequences. Read counts were proportionally assigned to the corresponding gene or TE as annotated in the provided transposable element GTF file ‘GRCh38_GENCODE_rmsk_TE.gtf’ downloaded from: http://labshare.cshl.edu/shares/mhammelllab/www-data/Tetranscripts/TE_GTF/.

The independent filtering option in the subsequent differential DESeq2 analysis was switched off to not exclude lowly expressed genes. The fold change threshold was set to 2 and FDR threshold to 0.01.

Transcriptional factors (TF) activity analysis was performed using ISMARA^46^. Quality and adaptor trimmed RNA-seq files for all OCI-AML3 samples (DMSO and CM-272) were uploaded to https://ismara.unibas.ch/mara/ for processing, followed by sample average. As genome build hg38 was not supported at the time of analysis, hg19 was used for ISMARA analysis. To determine a directional z-score for each enriched TF motif identified, the z-score for each given motif was multiplied by the sign of the Pearson correlation between each motif and its target genes and the direction of change in expression for said target genes (i.e., −1 for down-regulated genes and +1 for up-regulated genes).

### DNA methylation profiling and downstream analysis

DNA samples isolated from OCI-AML3 cells (DMSO and CM-272) were analyzed for global DNA methylation status using the Methylflash Methylated DNA Quantification Kit (Epigentek Group Inc) following the manufacturer’s instructions and measured in the FLUOstar Omega microplate reader.

Assessed DNA samples were bisulfite-converted using an EZ DNA methylation kit (Zymo Research) and hybridized onto an Infinium® MethylationEPIC BeadChip array (Illumina). IDAT files which were used as raw intensity data for methylation analysis using RnBeads 2.0 R package^102^ with a total 866.895 sites before filtering. First, probes giving unreliable results, determined by Greedycut algorithm (n=1.136), were filtered out. Additionally, cross reactive probes (n=44.726) and probes whose last 5 bases in its target sequence overlap with SNPs that were previously identified^103^, (n=19.828) were removed. The remaining data were normalized using beta-mixture quantile normalization (BMIQ)^104^ and background subtraction. Differentially methylated CpGs (DMCpGs) were identified based on differences in DNA methylation means (β-values) between CM-272 and DMSO-treated samples (Δβ ≥ 0.05 and p < 0.05).

Functional annotation enrichment analysis was performed using GREAT tool v4.0.4 (http://great.stanford.edu/public/html) ^105,106^ by mapping differentially methylated CpG (DMCpG) sites to the single nearest gene. All CpG probes which were found in the EPIC array were used as background in this analysis. Significantly enriched gene ontology biological process categories with p-value of <0.01 were used.

Using Homer v4.11 software^58^, 500 bp upstream and downstream of each DMCpG were assessed to predict transcription factors’ binding motifs. Intervals of 1000 bp of all CpG probes which are found in EPIC array were used as background in this analysis. Only known-motif enrichments were considered.

Chromatin state enrichment analysis was performed using EpiAnnotator tool^107^. DMCpGs were annotated with 15-Chromatin states using ChromHMM data set of reference human leukemia cell line K562 from the UCSC Genome Browser (https://genome.ucsc.edu/). Annotations of all CpG probes which are found on the EPIC array were used as background in this analysis.

### Software and Statistical Methods

R version 4.1.2 and Rstudio version 2023.03.0+386 Cherry Blossom^108^ were used for gene clustering analysis of the DNMT/HMT and heatmap rendering^109^ of the RNA-seq datasets TCGA-LAML and BeatAML downloaded from the cBioPortal for Cancer Genomics (http://cbioportal.org). Statistical evaluation and graphical reports were established with Graphpad Prism version 9.2.0 (GraphPad Software, San Diego, California, USA). Two-tailed Pearson’s correlation test (Spearman R) and one-way ANOVA with Dunnett’s correction for multiple comparisons (each condition vs. healthy bone marrow) were used for in silico RNA-seq AML datasets analysis. The Kaplan-Meier Log-Rank (Mantel-Cox) test was used for survival analysis. Unpaired Welch’s t-test and one-way ANOVA was used to assess significance in in vitro and in vivo assays. All in vitro experiments were performed in at least three independent experiments, each performed in three technical replicates. P values < 0.05 were considered significant with *, P < 0.05; **, P < 0.01; ***, P <0.001 and ****, P <0.0001. Adobe Illustrator version 27.8 and Biorender (BioRender.com) were used for graphics.

## Data availability

The RNA-Seq data sets generated and analyzed during the current study are available in the GEO repository as a Super Series under Accession ID # GSE241669.

## Acknowledgments

The authors would like to thank Dr. Ernesto Bockamp for their critical review of the manuscript; and Franziska Wille for their technical assistance and scientific advice. Support by the IMB Genomics Core Facility (Mainz) and the use of its NextSeq500 (funded by the Deutsche Forschungsgemeinschaft (DFG, German Research Foundation – 329045328) is gratefully acknowledged. For the DNA methylation array, the authors would like to thank the Genomics & Proteomics Core Facility (GPCF) of DKFZ at Heidelberg. For tissue processing, the authors would like to thank the pathology core facility of the following biobank locations: MTBIO (Munich), BBM (Mainz) and CHOICE (Dresden/Leipzig).

## Funding sources

This study was supported in part by grants from Deutsche Jose Carreras Leukemia Stiftung (DJCLS_04 R-2018) to B.G.; joint-funding from the German Consortium for Translational Cancer Research (DKTK CHOICE consortium) to B.G., K.G., U.P. and H.M. Work was also supported in part by funding from the Deutsche Forschungsgemeinschaft (DFG FOR2674 subprojects A1, A9 to C.P. and DFG, Project ID 318346496 – SFB 1292 to M.T.).

## Author contributions

B.G. designed the research, analyzed and interpreted the data, wrote and revised the manuscript, and supervised the study; A.S. and C.Z. performed the experiments, analyzed and interpreted the data, wrote the original draft of the manuscript and edited the manuscript; E.S., K.W., A.W., V.S., S.T., L.F. and A.S. performed experiments, analyzed and interpreted the data; C.W. generated and produced OCI-AML3-Luc-GFP line, F.R, S.K, C.A, C.G. and P.L, performed bioinformatics analyses and interpreted the data; D.S., T.K., M.T., J.R., J.S.H., K.S.G and U.P. provided administrative support and patient samples and revised the manuscript; C.G., J.P., C.P. and H.M. analyzed and interpreted the data and revised the manuscript.

## Competing interests

The authors declare no potential conflicts of interest.

## Notes

### Competing Interest Statement

The authors have declared no competing interest.

