## Supplemental data for "DNMT/G9a Complex Inhibition Uncovers Epigenetic Vulnerabilities and Induces IFN-Response in Acute Myeloid Leukemia"

### Supplementary Table 1 - Clinical data of Bone Marrow Biopsies

| ID | Age | Sex | Diagnosis | % Blasts | Karyotype | NPM1mut | FLT3mut | Other mutations |
|---|---|---|---|---|---|---|---|---|
| Healthy#1 | 63 | f | Control | NA | NA | WT | WT | WT |
| Healthy#2 | 68 | f | Control | NA | NA | WT | WT | WT |
| Healthy#3 | 67 | f | Control | NA | NA | WT | WT | WT |
| Healthy#4 | 67 | f | Control | NA | NA | WT | WT | WT |
| Healthy#5 | 63 | f | Control | NA | NA | WT | WT | WT |
| Healthy#6 | 72 | f | Control | NA | NA | WT | WT | WT |
| Healthy#7 | 66 | m | Control | NA | NA | WT | WT | WT |
| Healthy#8 | 79 | f | Control | NA | NA | WT | WT | WT |
| Healthy#9 | 66 | f | Control | NA | NA | WT | WT | WT |
| Healthy#10 | 82 | f | Control | NA | NA | WT | WT | WT |
| AML#1 | 64 | f | AML | 20-40% | 46, XX | MUT | WT | none |
| AML#2 | 59 | f | AML | 90% | unknown | MUT | ITD+ (ratio 0,513) | none |
| AML#3 | 77 | f | AML | 75% | 46 XX | MUT | ITD+ (ratio 0,237) | none |
| AML#4 | 82 | f | AML | 70% | 46 XX | MUT | ITD+ | none |
| AML#5 | 44 | m | AML | unknown | 46,XY,t(6;6)(p21;q21)[2] | MUT | WT | unknown |
| AML#6 | 76 | m | AML | 25-30% | 46,XY | MUT | TKD (ratio: 0,350) | none |
| AML#7 | 57 | m | AML | 80% | unknown | MUT | ITD+ | none |
| AML#8 | 52 | f | AML | 90% | 46,XX | MUT | ITD+ (ratio: 1,579) | none |
| AML#9 | 69 | m | AML | 30% | 46,XY | MUT | ITD+ (ratio 0,713) | none |
| AML#10 | 72 | m | AML | 90% | 46,XY | MUT | ITD+ | none |
| AML#11 | 67 | m | AML | 90% | 46,XY | MUT | WT | none |
| AML#12 | 46 | m | AML | 40% | 45, X,-Y | MUT | WT | none |
| AML#13 | 74 | f | AML | 40% | 46,XX | MUT | WT | none |
| AML#14 | 54 | f | AML | 90% | 46,XX | MUT | WT | none |
| AML#15 | 63 | f | AML | unknown | 46,XX | MUT | WT | none |

**Supplementary Table 2 - Clinical data of AML samples**

| UPN# | Type | Material | % Blast | Key mutations | Karyotype (G-Banding) |
|---|---|---|---|---|---|
| **Pt#1** | *secondary AML* | Bone marrow | *48%* | *FLT3-ITD; NPM1c; DNMT3a-R882* | 46,XX[4] |
| **Pt#2** | *de Novo AML (Early diagnosis)* | Bone marrow | *77%* | *FLT3-ITD; NPM1c* | 50 XY,+5,+6,+8,+19[7]/46, XY |
| **Pt#3** | *de Novo AML (Early diagnosis)* | PBMC | 75% | *FLT3-ITD; NPM1c* | 46,XX[20] |
| **Pt#4** | *secondary AML* | PBMC | 64% | *FLT3-ITD; NPM1c; DNMT3a-R882* | 46,XX[9] |
| **Pt#5** | *de Novo AML (Early diagnosis)* | Bone marrow | 72% | *FLT3-ITD; NPM1c* | 46,XY[2] |

### SUPPLEMENTARY FIGURE 1

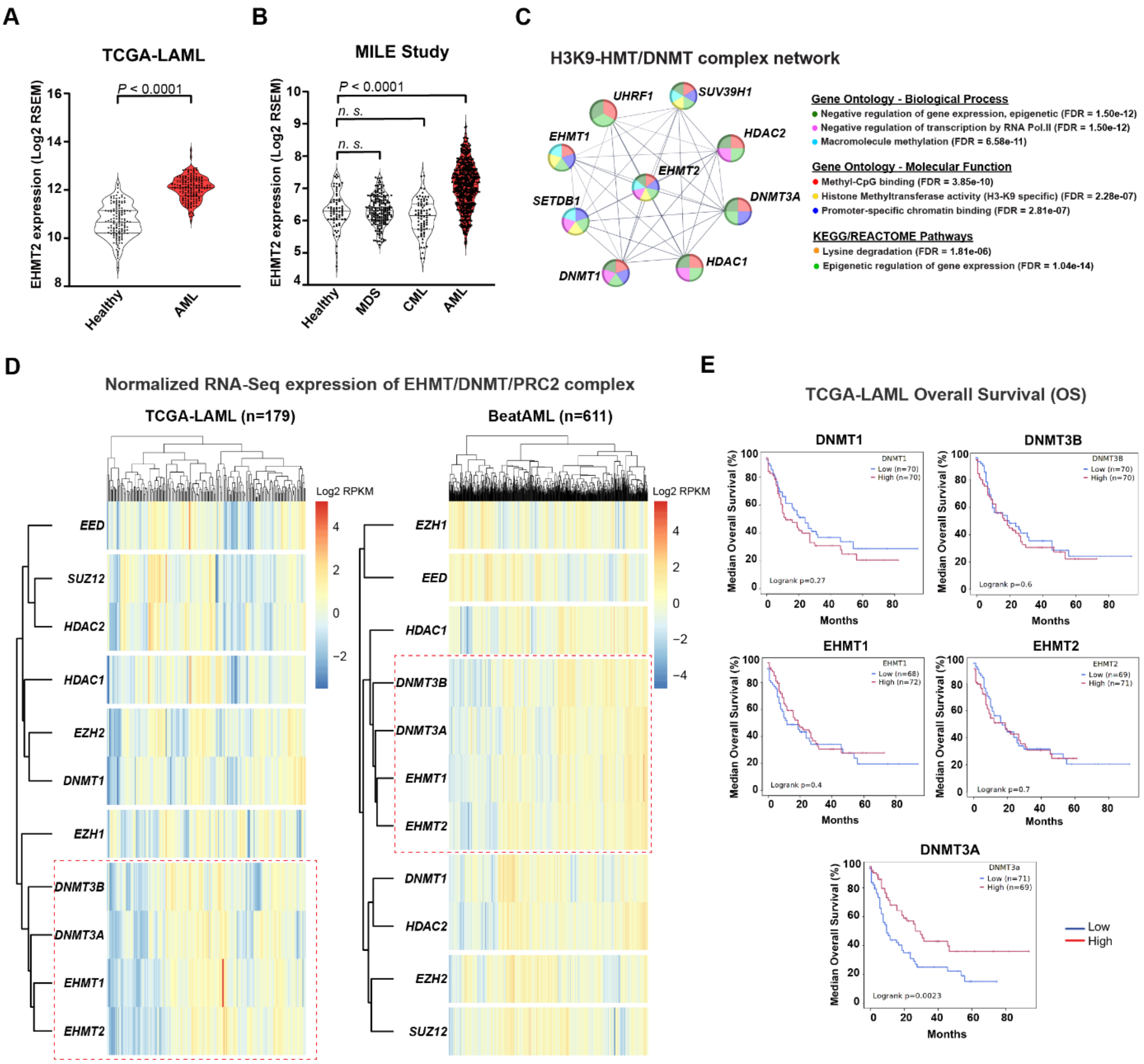


**Supplementary Figure 1. Correlation of G9a co-expression with HMT/DNMT family complex in AML and their association with patient survival.**

**(A-B)** *EHMT2 (G9a)* mRNA expression (normalized RNA-seq reads) between Healthy and myeloid diseases from **(A)** the TCGA-LAML study (n=179) **(B)** and the MILE study (n=542 samples). Within each Violin plot, a circle represents one patient value, dotted lines delimit first and third quartiles and black bars indicate the median. *P*-value statistical significance was assessed by one-way ANOVA with Dunnett's correction for multiple comparisons (each condition vs. healthy bone marrow). The non-indicated comparisons were not significant ($P > 0.05$). **(C)** Mapping view of relevant functional association network of members of the H3K9-HMT/DNMT complex generated using STRING protein interaction database; stronger associations are represented by thick lines. Each color correspond to significant epigenetic pathways associated with involved genes along with their respective adjustment for false discovery rate (FDR). **(D)** Hierarchical cluster analysis of TCGA-LAML and BeatAML studies for the genes involved in the H3K9-HMT/DNMT/PRC2 complex (11 genes). **(E)** Kaplan–Meier curve representing overall survival analysis of AML patients from the TCGA-LAML study ($n$ = 148) for individual mRNA expression of EHMT1, EHMT2, DNMT1, DNMT3A and DNMT3B above and below the median (as indicated in the graph). Statistical significance for *P*-value and hazard ratio (HR) was assessed by log-rank and Cox test.

### SUPPLEMENTARY FIGURE 2

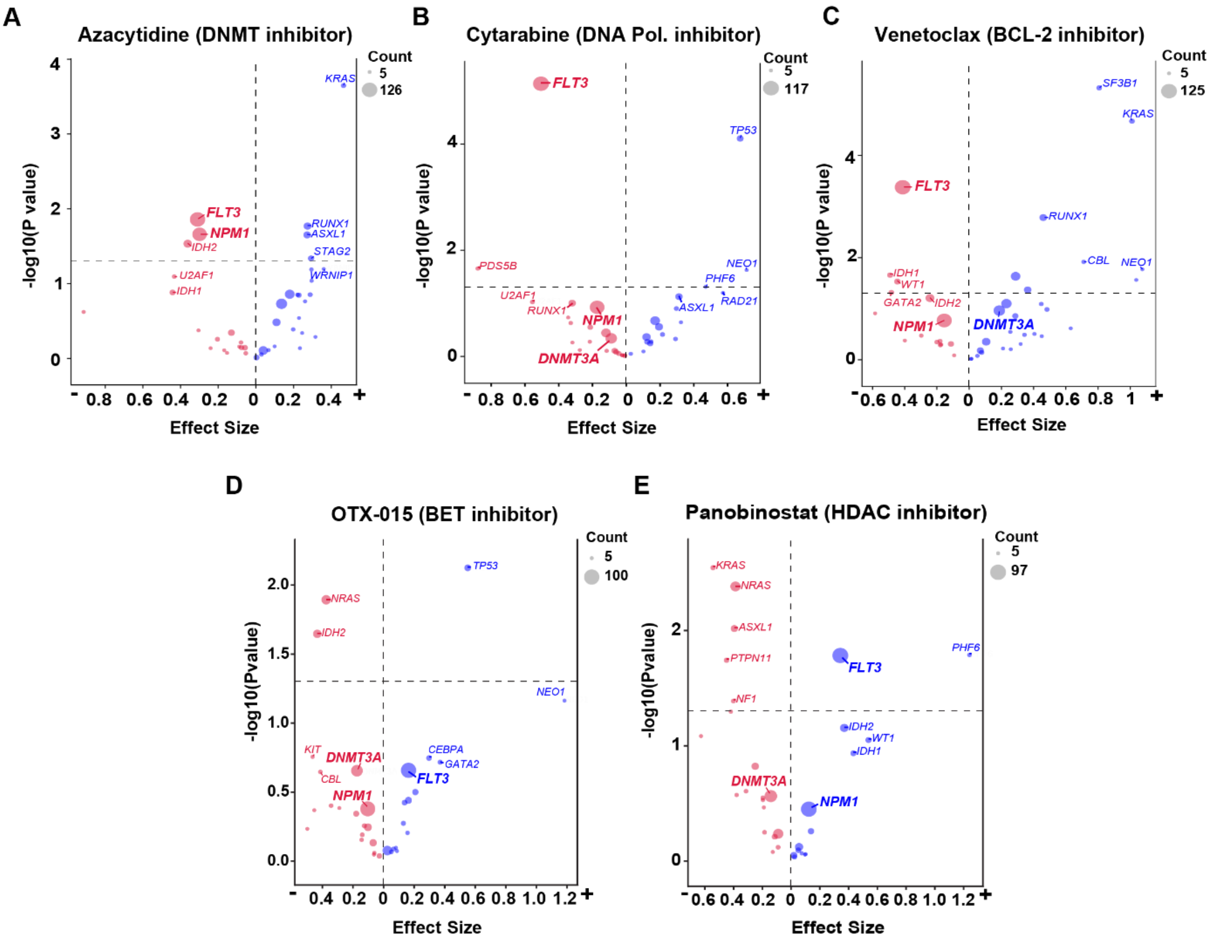


**Supplementary Figure 2. Drug sensitivity according to AML mutations.**

Visualizations of ex-vivo drug sensitivity by volcano plots according to AML subtypes from the BeatAML study towards **(A)** Azacytdine, **(B)** Cytarabine, **(C)** Venetoclax, **(D)** OTX-015 **(E)** and Panobinostat. Each circle correspond to a specific AML mutation composed of a minimum of 5 tested samples. Increased sensitivity is indicated by red and increased resistance indicated by blue as determined by the sign of the Glass's Delta effect size (X-axis) and dotted lines indicate limit of significance. All data analysis were retrieved and downloaded from the portal http://www.vizome.org/aml2/inhibitor/

### SUPPLEMENTARY FIGURE 3

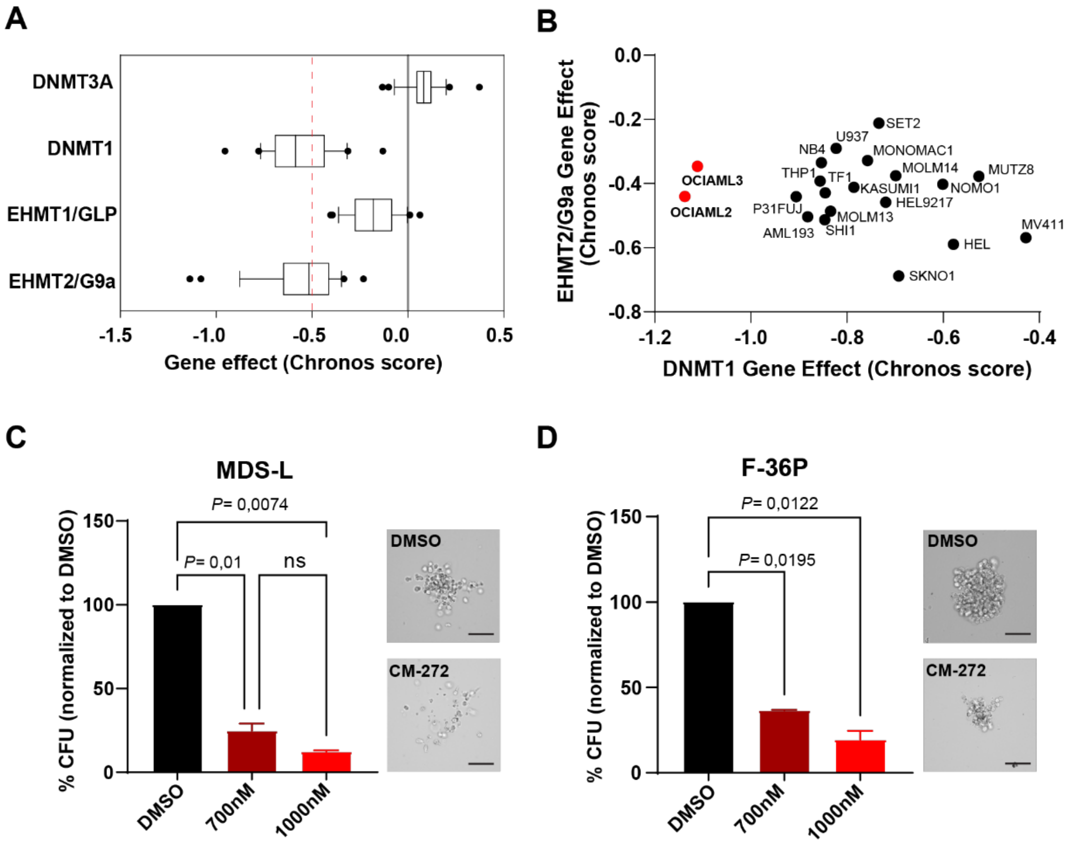


**Supplementary Figure 3. Dependency of AML cell lines toward DNMT/G9a.**

**(A-B)** DepMap analysis of the dependency of AML cell lines in CRISPR databases (CCLE Broad Institute). All data analysis were retrieved and downloaded from the portal https://depmap.org/portal/ **(A)** Dependency score of AML cell lines toward the individual CRISPR deletion of H3K9-HMT and DNMT genes. Whisker plots (with s.d) shows the distribution for each gene across all AML lines. Dots represents outliers and dotted red line indicate limit of significance for CRISPR gene effect (chronos score). **(B)** Dependency score of AML cell lines toward the dual deletion of DNMT1 and G9a genes. Each dot represent an AML cell line and red dots indicate the most sensitive AML cell lines according to CRISPR gene effect (chronos score). **(C-D)** Colonies outgrowth of MDSL **(C)** and **(D)** F-36P cells after CM-272 treatment at different indicated concentrations. CFU assays were carried out in technical duplicates and scored after 10-12 days incubation in methylcellulose. Data are presented as colonies percentage normalized to DMSO control ± s.d. One-way ANOVA was performed for *p*-value statistical. Micrographs of representative CFUs and Giemsa-stained cytospins were taken at x60 amplification.

### SUPPLEMENTARY FIGURE 4

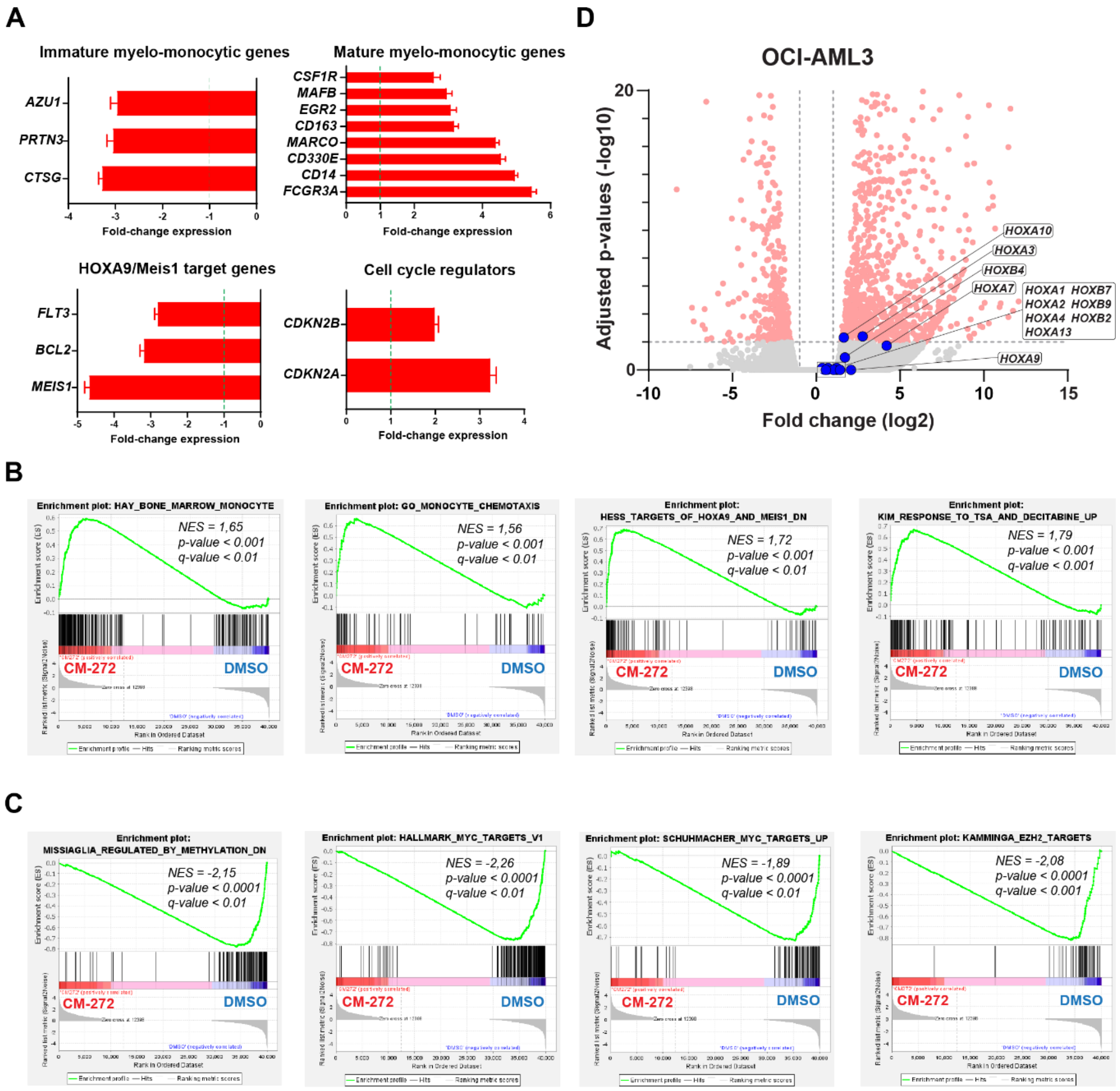


**Supplementary Figure 4. Changes in gene expression and related pathways in OC-AML3 upon treatment with CM-272.**

**(A)** Most significant genes up- and down-regulated in OCI-AML3 after CM-272 treatment. Bar graph represents fold change expression [mean ± standard deviation (s.d)] of RNA-seq and dotted line indicate limit of significance toward DMSO. **(B-C)** GSEA plot analysis showing the most representative gene signatures that are **(B)** up-regulated and **(C)** down-regulated in OCI-AML3 post-CM272 treatment. **(D)** Magnification of Volcano plots of OCI-AML3 RNA-seq treated with CM-272 (200nM). HOX genes are labelled in blue and horizontal dotted line indicate limit of statistical significance toward background.

### SUPPLEMENTARY FIGURE 5

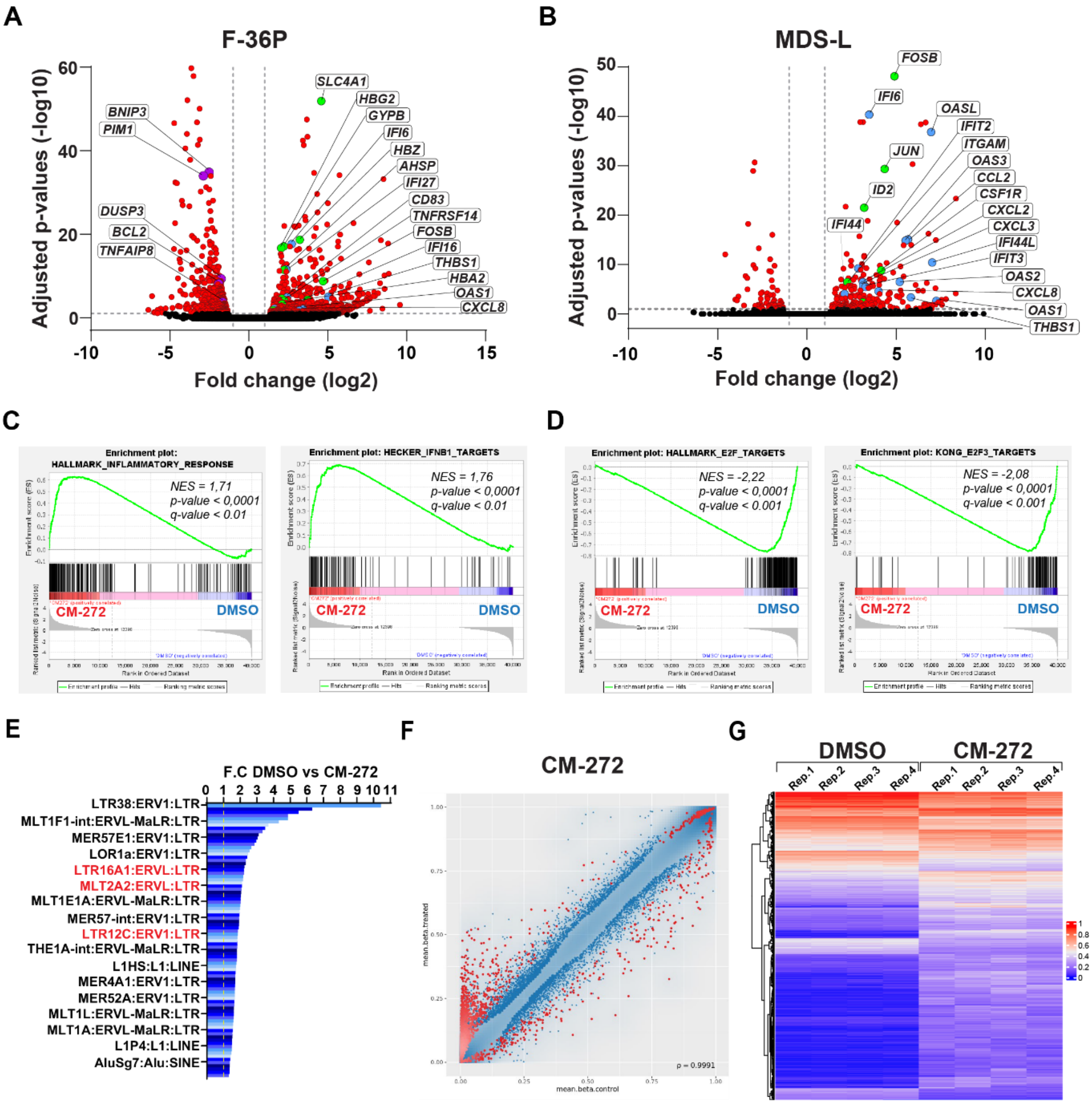


**Supplementary Figure 5. Gene expression and DNA methylation changes upon treatment with CM-272.**

**(A-B)** Volcano plots of RNA-seq data obtained from **(A)** F-36P and **(B)** MDS-L treated with CM-272 (700nM and 1000nM respectively). Genes involved in erythro-myeloid differentiation, interferon response and apoptosis are labeled in green, blue and magenta respectively. **(C-D)** GSEA plot analysis in OCI-AML3 post-CM272 treatment showing enrichment for **(C)** Interferon response-related gene signatures and **(D)** E2F-related gene signatures. **(E)** Most representative up-regulated dsRNAs of LTR/ERV (85) from RNA-seq of OCI-AML3 treated with CM-272. ERVs associated with antiviral immune responses are indicated in red font. Dotted yellow line represent limit of significance in comparison to DMSO. **(F)** Density plots of mean methylation beta values comparing control- and CM272-treated OCI-AML3 cells (n=4 each condition) using Infinium MethylationEPIC arrays. Point density is indicated by blue shading and corresponding Pearson's correlation value. Red dots represent single CpGs which are significantly hypo- or hypermethylated (FDR-adjusted p-value less than 0.05). **(G)** Hierarchical cluster analysis of global DMCpGs comparing control- and CM272-treated OCI-AML3 cells (n=4 each condition). Hyer- and Hypo-methylated DMCpGs are colored in red and blue respectively.

### SUPPLEMENTARY FIGURE 6

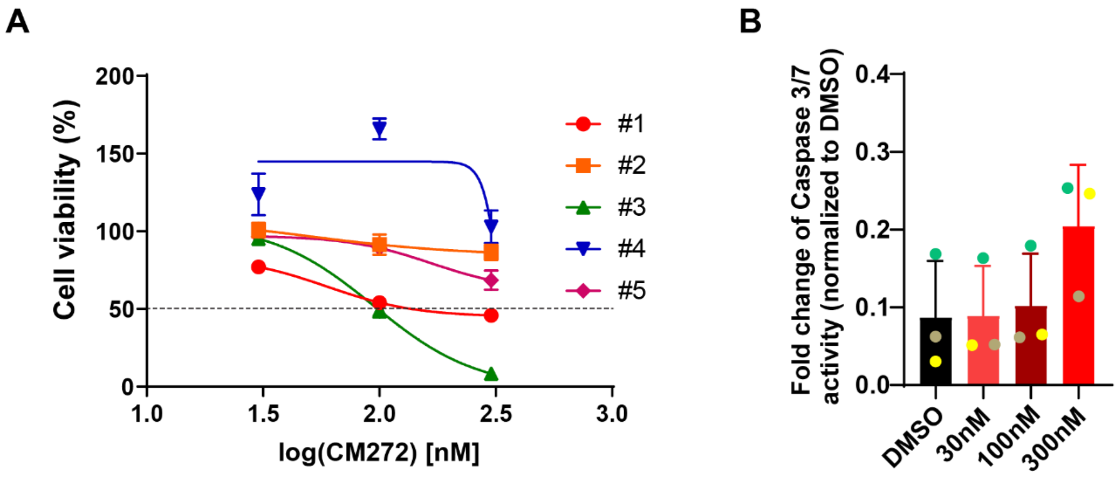


**Supplementary Figure 6. Dose response and cytotoxicity of patient-derived NPM1c AML blast cells upon treatment with CM-272.**

**(A)** Dose-response curves from cell-viability assays of NPM1c AML blast cells after 72h treatment with CM-272. Each line/curve represent an individual patient-derived NPM1c sample. Drug treatments were normalized to DMSO values and optimal dose range were determined from three independent experiments performed in technical triplicate. **(B)** Cytotoxicity analysis with a concentration range of CM-272 (single dose, 72h). Bar graph represents fold change of apoptosis measured by caspase3/7 expession [mean ± standard deviation (s.d)] and normalized to DMSO. Each color dot represents an individual patient-derived NPM1c AML sample (3 patients in total).
